# A genomic assessment of the marine-speciation paradox within the toothed whale superfamily Delphinoidea

**DOI:** 10.1101/2020.10.23.352286

**Authors:** Michael V Westbury, Andrea A. Cabrera, Alba Rey-Iglesia, Binia De Cahsan, David A. Duchêne, Stefanie Hartmann, Eline D Lorenzen

## Abstract

The importance of post-divergence gene flow in speciation has been documented across a range of taxa in recent years, and may have been especially widespread in highly mobile, wide-ranging marine species, such as cetaceans. Here, we studied individual genomes from nine species across the three families of the toothed whale superfamily Delphinoidea (Delphinidae, Phocoenidae, Monodontidae). To investigate the role of post-divergence gene flow in the speciation process, we used a multifaceted approach, including: (i) phylogenomics, (ii) the distribution of shared derived alleles, and (iii) demographic inference. We found the divergence of lineages within Delphinoidea did not follow a process of pure bifurcation, but was much more complex. Sliding-window phylogenomics reveal a high prevalence of discordant topologies within the superfamily, with further analyses indicating these discordances arose due to both incomplete lineage sorting and gene flow. D-statistics, D-foil, and *f*-branch analyses supported gene flow between members of Delphinoidea, with the vast majority of gene flow occurring as ancient interfamilial events. Demographic analyses provided evidence that introgressive gene flow has likely ceased between all species pairs tested, despite reports of contemporary interspecific hybrids. Our study provides the first steps towards resolving the large complexity of speciation within Delphinoidea; we reveal the prevalence of ancient interfamilial gene flow events prior to the diversification of each family, and suggests that contemporary hybridisation events may be disadvantageous, as hybrid individuals do not appear to contribute to the parental species’ gene pools.

## Introduction

The formation of new species involves the divergence of lineages through reproductive isolation. Isolation can initially occur in allopatry (geographical isolation without gene flow) or in sympatry (biological/ecological isolation with gene flow). Over time, isolation can be maintained and strengthened, ultimately leading to the formation of new species (Norris and Hull, 2012). While allopatric speciation requires geographical isolation plus time, sympatric speciation often requires a broader and more complicated set of mechanisms (Turelli, Barton and Coyne, 2001). These mechanisms mostly rely on ecologically mediated natural selection. Parapatric speciation, on the other hand, encompasses intermediate scenarios of partial, but incomplete, physical restrictions to gene flow leading to speciation.

Through the analysis of whole-genome datasets, the detection of post-divergence gene flow in various distinct taxonomic groups is becoming commonplace (Árnason *et al*., 2018; Barlow *et al*., 2018; Westbury *et al*., 2020), demonstrating that speciation is much more complex than a simple bifurcating process (Feder, Egan and Nosil, 2012; Campbell and Poelstra, 2018). Speciation is not an instantaneous process, but usually requires tens of thousands to millions of generations to achieve complete reproductive isolation (Liu *et al*., 2014; Butlin and Smadja, 2018). The duration it takes to reach this isolation may be especially long in highly mobile marine species, such as cetaceans, due to a relative lack of geographic barriers in the marine realm, and therefore high potential for secondary contact and gene flow (Árnason *et al*., 2018).

The apparent inability to undergo allopatric speciation in marine species has been termed the marine-speciation paradox (Bierne, Bonhomme and David, 2003). However, over the past decade, genomic studies have provided insights into how speciation can occur within cetaceans (Árnason *et al*., 2018; Moura *et al*., 2020). For example, initial phases of allopatry among populations of killer whales (*Orcinus orca*) may have led to the accumulation of ecological differences between populations, which strengthened population differences even after secondary contact (Foote *et al*., 2011; Foote and Morin, 2015). However, whether these initial phases of allopatry caused the divergence, or whether speciation occurred purely in sympatry, remains debated (Moura *et al*., 2015; Foote, 2018). But, these two hypotheses are not necessarily mutually exclusive. Instead, differentiation in parapatry, encompassing features of both allopatric and sympatric speciation, may have been key in the evolutionary history of cetaceans.

The toothed whale superfamily Delphinoidea represents an interesting opportunity to further explore speciation in the presence of putative interspecific gene flow. The crown root of Delphinoidea has been dated at ^~^19 million years ago (Ma) (95% CI 19.73 - 18.26 Ma) (McGowen *et al*., 2020) and has given rise to three families: (i) Delphinidae, the most species-rich family, which comprises dolphins and ‘black-fish’ (such as killer whales and pilot whales (*Globicephala spp*.)); (ii) Phocoenidae, commonly known as porpoises; and (iii) Monodontidae, which comprises two extant lineages, beluga (*Delphinapterus leucas*) and narwhal (*Monodon monoceros*).

Delphinoidea is of particular interest, as contemporary interspecific hybrids have been reported within all three families (Delphinidae (Miyazaki *et al*., 1992; Silva, Silva and Sazima, 2005; Espada *et al*., 2019); Phocoenidae (Willis *et al*., 2004); Monodontidae (Skovrind *et al*., 2019)). However, these represent recent hybridization events that occurred long after species divergence, and their contribution to the parental gene pools is mostly unknown. The presence of more ancient introgressive hybridization events between families, and during the early radiations of these families, has yet to be investigated. With the rapid increase of genomic resources for cetaceans, and in particular for species within Delphinoidea, we are presented with the ideal opportunity to investigate post-divergence gene flow between lineages, furthering our understanding of speciation processes in cetaceans.

Here, we utilise publicly available whole-genome data from nine species of Delphinoidea, representing all three families, to investigate signs of post-divergence gene flow across their genomes. Our analyses included five Delphinidae (killer whale, Pacific white-sided dolphin (*Lagenorhynchus obliquidens*), long-finned pilot whale (*Globicephala melas*), bottlenose dolphin (*Tursiops truncatus*), Indo-Pacific bottlenose dolphin (*T. aduncus*)); two Phocoenidae (harbour porpoise (*Phocoena phocoena*), finless porpoise (*Neophocaena phocaenoides*)); and two Monodontidae (beluga, narwhal). Moreover, we compare their species-specific genetic diversity and demographic histories, and explore how species abundances may have played a role in interspecific hybridisation over the last two million years.

## Results and discussion

### Sliding window phylogenomic analyses

To assess the evolutionary relationships across the genomes of the nine Delphinoidea species investigated, we computed non-overlapping, sliding-window, maximum-likelihood phylogenies of four different window sizes in RAxML (Stamatakis, 2014). These analyses resulted in 43,207 trees (50 kilobase (kb) windows), 21,387 trees (100 kb windows), 3,705 trees (500 kb windows), and 1,541 trees (1 megabase (Mb) windows) (Fig. 1, Supplementary Fig. S1, Supplementary Table S1). The 50 kb windows retrieved 96 unique topologies, 100 kb windows retrieved 47 unique topologies, 500 kb windows retrieved 16 unique topologies, and 1 Mb windows retrieved 15 unique topologies. Regardless of window size, we retrieved consensus support for the species tree previously reported using target-sequence capture (McGowen *et al*., 2020). However, when considering the smallest window size (50 kb), we found a considerable proportion of trees (up to 76%) with an alternative topology to the species tree (Fig. 1A). These alternative topologies may be due to incomplete lineage sorting (ILS) or interspecific gene flow (Leaché *et al*., 2014). Moreover, the higher prevalence of this pattern in the shorter 50 kb windows may indicate that inconsistencies in topology are caused by ancient, rather than recent, gene flow events, as recombination is expected to break up longer introgressed regions over time (as a comparison, only 21% of windows in the 1 Mb dataset do not show the most common topology, Fig. 1B).

**Figure 1:**
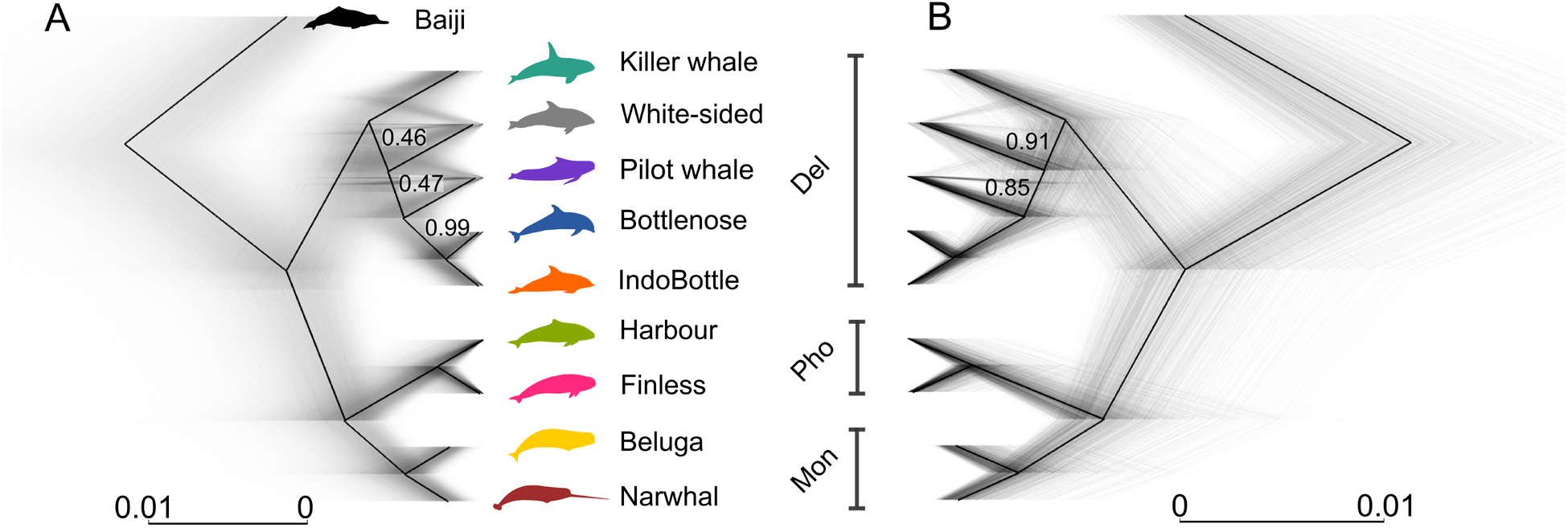
Sliding-Window Maximum likelihood trees of nine Delphinoidea species and the baiji. The trees were constructed using non-overlapping sliding windows of (A) 50 kb in length and (B) 1 Mb in length. Black lines show the multi-species coalescent species tree estimate, grey lines show individual trees. Numbers on branches show the proportion of windows supporting the node. Branches without numbers had maximal support. Bottlenose dolphin silhouette: license Public Domain Dedication 1.0; remaining Delphinoidea silhouettes: Chris huh, license CC-BY-SA-3.0 (https://creativecommons.org/licenses/by-sa/3.0/).

We explored whether the large number of phylogenetic discrepancies in the 50kb windows could be linked to the GC content (%GC) of the windows as elevated levels of GC content can result from higher levels of GC-Biased Gene Conversion (gBGC) in regions with higher levels of recombination (Lartillot, 2013). When binning windows into either high, medium, or low levels of GC content, the most common topologies were consistent, but with slight differences in overall values (Supplementary Table S2). This result suggests that the topological discrepancies are not arising purely due to GC-content linked biases and recombination rate.

### Separating ILS and gene flow

To investigate whether the alternative topologies could simply be explained by ILS, or a combination of ILS and gene flow, we ran Quantifying Introgression via Branch Lengths (QuIBL) (Edelman *et al*., 2019) on every twentieth tree from the 50 kb sliding-window analysis (Supplementary Table S3), as well as on a dataset that contained trees constructed using 20 kb windows with a 1 Mb slide (Supplementary Table S4). We were only able to investigate the potential cause of discordances within the Delphinidae family, as we did not recover any phylogenetic discordances between families, and all families were respectively monophyletic.

When considering the results using 50 kb windows, we found significant evidence of ILS and gene flow in all species pairwise comparisons within Delphinidae. The only comparisons that did not show significant results for gene flow were those that contained both the bottlenose and Indo-Pacific bottlenose dolphins. The lacking evidence of gene flow when both *Tursiops* species were included, suggests signals of gene flow between either *Tursiops* species and killer whale, Pacific white-sided dolphin, or pilot whale are likely remnants of ancestral gene flow events between the ancestral *Tursiops* and the given comparative species.

Similar to the 50 kb windows, the 20 kb window analysis showed a large proportion of alternative topologies within Delphinidae likely arose due to both ILS gene flow. Again, we retrieved most non-significant results when both *Tursiops* species were included in the analysis. Moreover, although we found no evidence of gene flow between killer whale and pilot whale when either *Tursiops* was included as the triplet outgroup, we found evidence of gene flow when the Pacific white-sided dolphin was the triplet outgroup. We also found no evidence for gene flow between the Indo-Pacific bottlenose and Pacific white-sided dolphins, regardless of triplet outgroup. It is difficult to ascertain why we observe discrepancies between results based on the triplet outgroup. But, taken together, our QuIBL analyses suggest a combination of ILS and gene flow played a role in shaping the evolutionary history of Delphinidae.

### Accounting for ILS in gene flow estimates

To further explore potential gene flow while taking ILS into account, we used D-statistics (Green *et al*., 2010; Durand *et al*., 2011). D-statistics uses a four-taxon approach [[[H1, H2], H3], Outgroup] to uncover the differential distribution of shared derived alleles, which may represent gene flow between either H1/H3 or H2/H3. Here we used baiji (*Lipotes vexillifer*) as the outgroup, and alternated ingroup positions based on the consensus topology. In congruence with the QuIBL results, we found significant levels of gene flow within Delphinidae. However, we also found higher levels of gene flow between the killer whale, pilot whale, and Pacific white-sided dolphin and the Indo-Pacific bottlenose dolphin, relative to the bottlenose dolphin. In fact, 85 out of 86 tests showed significant signs of gene flow both within and between families (Supplementary Table S5). The only comparison that did not return a significant result was [[[finless porpoise, harbour porpoise], narwhal], outgroup]. This does not necessarily mean there was no gene flow between these species, but could be caused by equal amounts of gene flow between both porpoise species and narwhal. Such abundant signs of gene flow suggests the evolutionary history of Delphinoidea was more complex than a simple bifurcating process. Alternatively, our findings may reflect limitations of the D-statistic and false positives due to gene flow between ancestral lineages (Moodley *et al*., 2020).

### Direction of gene flow

Due to the inability of the four-taxon D-statistics approach to detect the direction of gene flow, as well as whether gene flow events may have occurred between ancestral lineages, we used D-foil (Pease and Hahn, 2015). D-foil enables further characterization of the D-statistics results, which may be particularly relevant given the complex array of gene flow putatively present within Delphinoidea. D-foil uses a five-taxon approach [[H1, H2] [H3, H4], Outgroup] and a system of four independent D-statistics in a sliding-window fashion, to uncover (i) putative gene flow events, (ii) donor and recipient lineages, and (iii) whether gene flow events occurred between a distantly related lineage and the ancestor of two sister lineages, which is indicative of ancestral-lineage gene flow. However, as the input topology requirements of D-foil are [[H1, H2] [H3, H4], Outgroup], we were only able to investigate gene flow between families, and not within families, using this analysis. Hence, we tested for gene flow between Delphinidae/Phocoenidae, Delphinidae/Monodontidae, and Phocoenidae/Monodontidae.

The D-foil results underscore the complex pattern of post-divergence gene flow between families indicated by the D-statistics. We found support for interfamilial gene flow events between all nine species investigated, to varying extents (Supplementary Table S6). This could reflect multiple episodes of gene flow between all investigated species. Alternatively, the pattern could reflect ancient gene flow events between the ancestors of H1-H2 and H3-H4 (in the topology [[H1, H2] [H3, H4], Outgroup]), with differential inheritance of the introgressed loci in subsequent lineages. Such ancestral gene flow events have previously been shown to lead to false positives between species pairs using D-statistics (Moodley *et al*., 2020). A further putative problem with these results can be seen when implementing D-foil on the topology [[Delphinidae, Delphinidae], [Monodontidae, Phocoenidae], Outgroup]. We found the majority of windows support a closer relationship between Delphinidae (ancestors of H1 and H2) and Monodontidae (H3), as opposed to the species tree. If this result is correct, it suggests the input topology was incorrect, and the results reflect more recent common ancestry and not gene flow. This would imply Delphinidae and Monodontidae are sister lineages, as opposed to Phocoenidae and Monodontidae. However, this falls in contrast with the family topology of [Delphinidae, [Phocoenidae, Monodontidae]] retrieved in our phylogenetic analyses under the multi-species coalescent (Fig. 1) and those reported by others (Steeman *et al*., 2009; McGowen *et al*., 2020).

Taken together, it is difficult to ascertain whether our D-statistics and D-foil results of prevalent gene flow among most species pairs are true, or whether some results may have arisen due to biases that can occur when attempting to infer gene flow between highly divergent lineages. False positives and potential biases in D-statistics and D-foil can arise due to a number of factors including (i) ancestral population structure, (ii) introgression from unsampled and/or extinct ghost lineages, (iii) differences in relative population size of lineages or in the timing of gene flow events, (iv) different evolutionary rates or sequencing errors between H1 and H2, and (v) gene flow between ancestral lineages (Slatkin and Pollack, 2008; Zheng and Janke, 2018; Moodley *et al*., 2020). These issues are important to consider when interpreting our results, as the deep divergences of lineages suggest the possibility for a number of ancestral gene flow events, as well as gene flow events between now-extinct lineages, that may bias results.

### Gene flow between ancestral lineages

Due to the large number of possible D-statistics comparisons, and difficulties disentangling false positives that may arise due to ancient gene flow events, we performed the *f*-branch test (Malinsky *et al*., 2018; Malinsky, Matschiner and Svardal, 2021). The test takes correlated allele sharing into account when visualising f4-ratio (similar to D-statistics) results. The *f*-branch results suggested several instances of gene flow, many between ancestral lineages with relatively small values of fb (<0.04 with the majority being ^~^0.01) (Fig. 2 and Supplementary Fig. S3). This result suggests widespread gene flow but in small quantities. However, it should be noted that fb represents relative quantities of gene flow and likely also decreases the older the introgression event (Martin, Davey and Jiggins, 2015)so the values we present here may not fully represent the absolute levels of gene flow. When considering interfamilial gene flow events, we see excess allele sharing (fb) between the ancestral Monodontidae branch and all Delphinidae species, which we interpret as gene flow between the ancestral lineages of Monodontidae and Delphinidae. We also uncovered elevated fb between the ancestor of all Delphinidae (to the exclusion of the killer whale) and all Phocoenidae and Monodontidae species, which could suggest gene flow between Delphinidae and the ancestral Phocoenidae/Monodontidae lineage. However, the exclusion of the killer whale may be due to the inability of the four taxon f4-ratio test to calculate gene flow between the killer whale and ancestral Phocoenidae/Monodontidae. Based on this limitation, we take a conservative approach and suggest this result reflects gene flow between the ancestral Delphinidae and ancestral Phocoenidae/Monodontidae (Fig. 2C).

**Figure 2:**
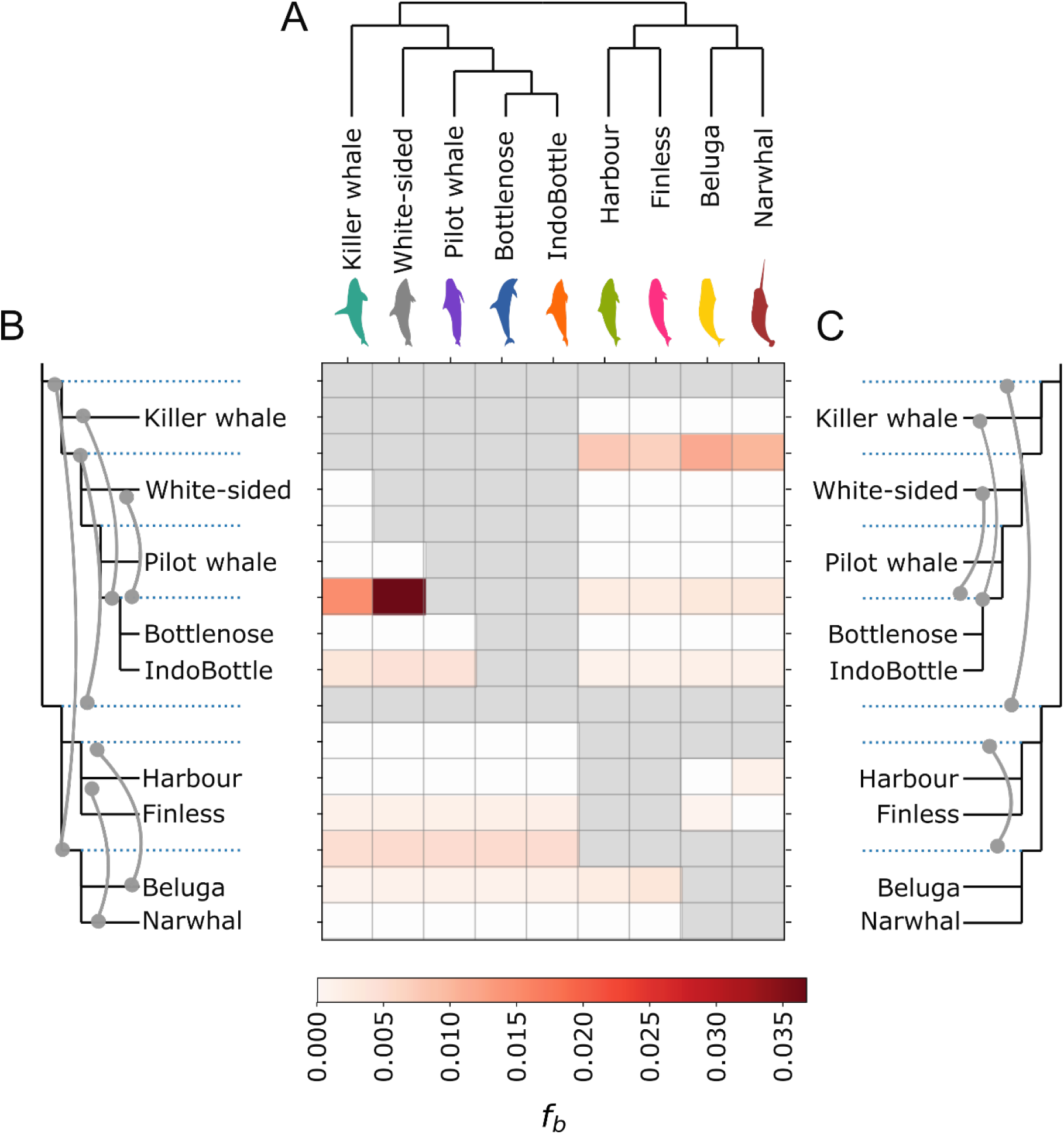
Genome-wide *f*-branch results. (A) Species tree; (B) and (C) Species tree in expanded form, with internal branches as dotted lines. The values in the matrix refer to excess allele sharing between the expanded tree branch (relative to its sister branch) and the species on the *x*-axis. Lines connecting branches show: (B) gene flow events inferred directly from the *f*-branch results; (C) gene flow events that we hypothesised from the *f*-branch results, while accounting for (i) the inability to detect gene flow between sister lineages, and (ii) a lack of a positive means less gene flow relative to the sister lineage, rather than no gene flow.

Further supporting the hypothesis of gene flow between the ancestral Delphinidae and ancestral Phocoenidae/Monodontidae (Fig. 2C), we also observed signs of gene flow between the finless porpoise and all Delphinidae species, which suggests gene flow between the finless porpoise and ancestral Delphinidae. This seems unreasonable, as the finless porpoise diverged from the harbour porpoise much more recently (^~^5 Ma) than the time to the most recent common ancestor (tMRCA) of all Delphinidae (^~^10 Ma, (McGowen *et al*., 2020), meaning gene flow would have occurred independently between the finless porpoise and almost every Delphinidae species studied here. Moreover, the *f*-branch showed similar fb between the Indo-Pacific bottlenose dolphin and all Phocoenidae and Monodontidae, as well as between the ancestral *Tursiops* and all Phocoenidae and Monodontidae. Similar to the finless porpoise and ancestral Delphinidae, this result seems unlikely due to the divergence times of *Tursiops*.

We also found signals of gene flow between beluga and both Phocoenidae species, but not between narwhal and Phocoenidae. This pattern may be more parsimoniously explained by an ancestral event between Phocoenidae and Monodontidae, where the narwhal retained less introgressed alleles. A given *f*b statistic presents the signal of excess gene flow relative to the ingroup’s sister taxa (Malinsky, Matschiner and Svardal, 2021). Hence, not recovering a signal of gene flow with the sister taxa does not mean it did not occur. Rather, gene flow may have occurred between taxa, but to a lesser degree. Taking this into account, we suggest our results may instead be remnants of ancestral gene flow events between the ancestral Phocoenidae and Monodontidae lineages (Fig. 2C). A lack of evidence for more recent, species-specific gene flow events here is congruent with the sliding-window and species tree analyses, which showed strong support for Phocoenidae and Monodontidae as sisters.

The *f*-branch test also revealed interspecific gene flow events within Delphinidae may have been common. We uncovered evidence for gene flow between the Pacific white-sided dolphin and ancestral *Tursiops*, as well as the killer whale and ancestral *Tursiops*. However, we are unable to dissect whether there was gene flow between the pilot whale and ancestral *Tursiops*, due to the limitation of the four-taxon requirement.

To investigate whether the X chromosome may have presented a more pronounced barrier to gene flow relative to the autosomes, we ran the *f*-branch test on scaffolds aligning to the X chromosome. Results were similar to the genome-wide dataset (Supplementary Figs. S2 and S4). The most obvious difference is that evidence for gene flow between Phocoenidae and Monodontidae is not as pronounced as in the genome-wide dataset. It is difficult to discern whether the lack of resolution here is due to the X chromosome constituting a smaller dataset, or whether parts of the X chromosome were not incorporated into the recipient gene pool due to the occurrence of more rapid reproductive isolation on the X chromosome (Payseur and Rieseberg, 2016). The former option appears more probable, due to the consistent evidence for gene flow between the beluga and both Phocoenidae species, which are likely the remnants of ancestral gene flow events between Phocoenidae and Monodontidae.

By combining results acquired through sliding-window phylogenies, QuIBL, D-statistics, Dfoil, and *f*-branch, we are able to better decipher the complex evolutionary history of Delphinidae, and the signatures of interspecific gene-flow events present in most individuals studied. We found the most probable explanation for such wide-spread signatures to be the differential inheritance of remnant loci from ancestral gene flow events. However, as exemplified here and due to the limitations of each method, uncovering the exact lineages involved in these events is challenging.

### Cessation of lineage sorting and/or gene flow

To further elucidate the complexity of interspecific gene flow within Delphinoidea, we implemented F1 hybrid PSMC (hPSMC) (Cahill *et al*., 2016) on the autosomes of our species of interest. This method creates a pseudo-diploid sequence by merging pseudo-haploid sequences from two different genomes, which in our case represents two different species. The variation in the interspecific pseudo-F1 hybrid genome cannot coalesce more recently than the emergence of reproductive isolation between the two parental species. If some regions within the genomes of two target species are yet to fully diverge, due to ILS or to gene flow, hybridisation may still be possible. Therefore, we can use this method to infer when reproductive isolation between two species may have occurred.

When considering the upper bound of when two target genomes coalesce (equating the oldest date), and the lower bound of each divergence date (equating the most recent date) (McGowen *et al*., 2020), we found the majority of comparisons (29/36) show lineage sorting and/or gene flow occurred for >50% of the post-divergence branch length (Fig. 3, Supplementary data - hPSMC). However, we used divergence times estimated assuming a fixed tree-like topology without taking gene tree discordances into account, which could lead to extended terminal branches and overestimated dates due to molecular substitutions of discordant loci needing to be placed somewhere on the tree (Mendes and Hahn, 2016). Nevertheless, our results suggest that reaching complete reproductive isolation in Delphinoidea was a slow process, due to ILS and/or gene flow. ILS levels are known to be proportional to ancestral population sizes, and inversely proportional to time between speciation events (Pamilo and Nei, 1988). Hence, if ILS was the only explanation for this phenomenon, this would suggest extremely large ancestral population sizes. We do indeed see that the species pairs with the highest Ne prior to the end of lineage sorting/gene flow (Supplementary table S7) also have the largest discrepancies between divergence date and the date at which the two genomes coalesce. However, an alternative, and perhaps more likely, explanation is the occurrence of gene flow after initial divergence, supported by our phylogenomic, D-statistics, Dfoil, and *f*-branch results above. Post-divergence gene flow may reflect the ability of cetacean species to travel long distances, and the absence of significant geographical barriers in the marine environment. Alternatively, if geographic barriers did drive initial divergence, the pattern retrieved in our data may reflect secondary contact prior to complete reproductive isolation.

**Figure 3:**
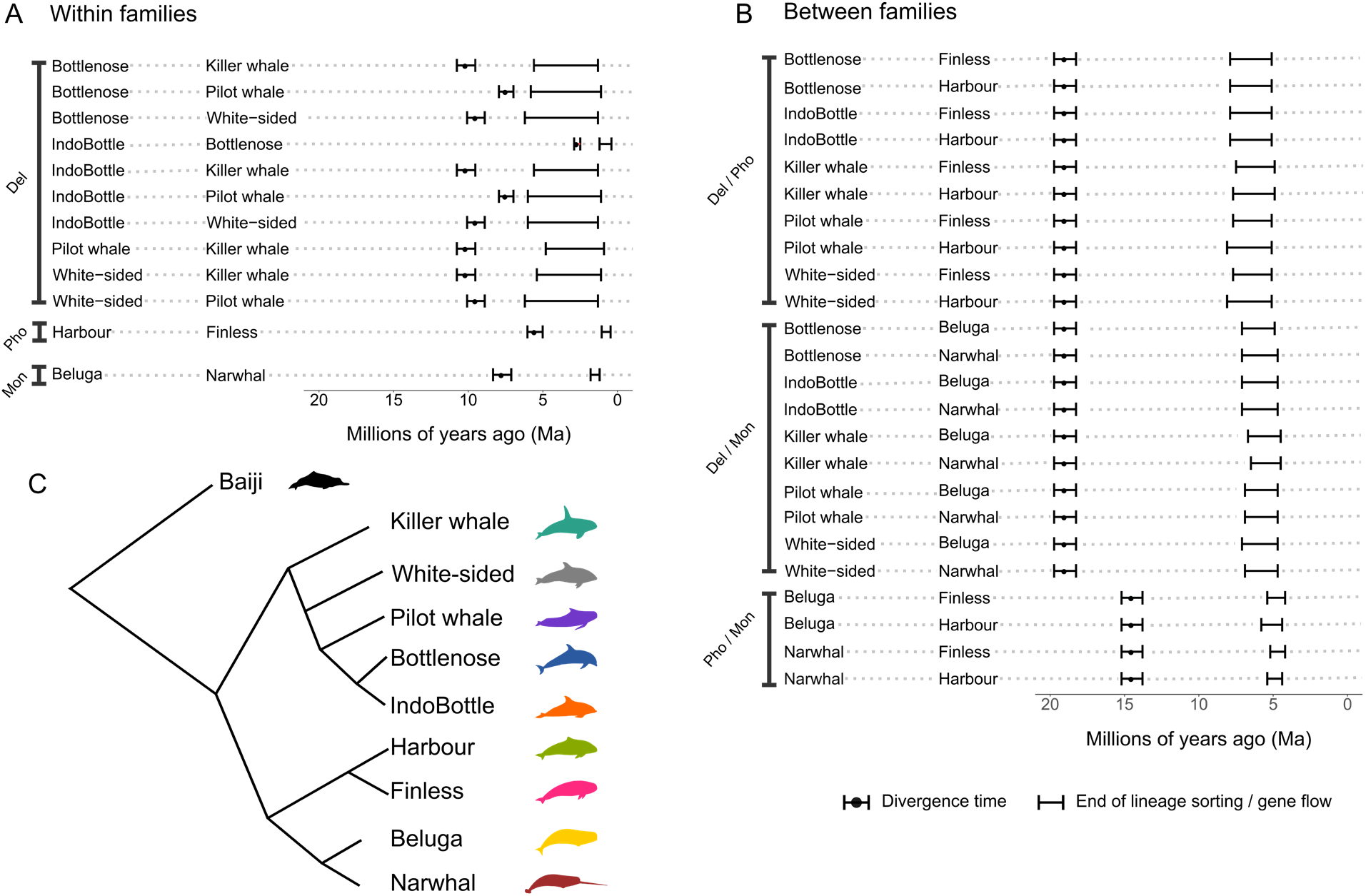
Estimated divergence times and time intervals during which gene flow ceased between species (A) within families and (B) between families. Estimated time intervals of when gene flow ceased between species pairs are based on hPSMC results. A PSMC analysis on a pseudo-F1 hybrid diploid genome between two species results in an asymptotic increase in Ne at the time point the two genomes coalesce. By simulating data with various timings of divergence, and finding the simulated data most closely matching the empirical data, we determined the time interval gene flow ceased (Supplementary results - hPSMC). Divergence time estimates are taken from McGowen *et al* 2020.

Our hPSMC results showed an almost simultaneous cessation of lineage sorting/gene flow regardless of species pair within the Delphinidae family (Fig 3A), as well as comparisons between families (Fig 3B). Based on our D-statistic/D-foil/*f*-branch results showing many of the signals of gene flow may be remnants of ancestral gene flow events, we hypothesise that our deep-time hPSMC results may also be produced by ILS of ancestrally introgressed regions. If we assume the divergence dates are correct, this hypothesis also offers an explanation regarding why the end of interfamilial ILS/gene flow occurs after the tMRCA of the family in many cases. For example, the tMRCA of Phocoenidae is ^~^6Ma, and the tMRCA of Monodontidae is ^~^7Ma but our hPSMC suggests that ILS/gene flow did not stop between Phocoenidae and Monodontidae until ^~^5Ma. Superficially, this implies that interfamilial gene flow occurred uniquely between beluga/finless porpoise, beluga/harbour porpoise, narwhal/finless porpoise, and narwhal/harbour porpoise, and ceased for all species pairs at the same time. While this may have been the case, a more likely explanation is that lineage sorting of introgressed regions from an ancestral gene flow event was not complete until the time periods that our hPSMC results recovered.

Despite our hPSMC results of long-term lineage sorting/gene flow in the majority of species comparisons, they also suggested that lineage sorting is complete and gene flow has ceased between all lineages in our dataset. This finding is in contrast with confirmed reports of fertile contemporary hybrids between several of our target species, and may reflect the inability of hPSMC to detect low levels of migration. For example, viable offspring have been reported between bottlenose dolphins and Indo-Pacific bottlenose dolphins (Gridley *et al*., 2018) and between bottlenose dolphins and Pacific white-sided dolphins (Miyazaki *et al*., 1992; Crossman, Taylor and Barrett-Lennard, 2016). Simulations have shown that in the presence of as few as 1/10,000 migrants per generation, hPSMC will suggest continued gene flow. However, this is not the case with a rate < 1/100,000 migrants per generation. Rather, in the latter case, the exponential increase in effective population size (Ne) of the pseudo-hybrid genome, which can be used to infer the date at which gene flow ceased between the parental species, becomes a more gradual transition, leading to a larger estimated time interval of gene flow (Cahill *et al*., 2016). Within Delphinidae, we observe a less pronounced increase in Ne in the pseudo-hybrids, suggesting continued, but very low migration rates (Supplementary results - hPSMC). This finding suggests that gene flow within Delphinidae may have continued for longer than shown by hPSMC, which may not be sensitive enough to detect low rates of recent gene flow. Either way, our hPSMC results within and between all three families showed a consistent pattern of long periods of lineage sorting/gene flow in Delphinoidea, some lasting more than ten million years post divergence.

We further assessed the robustness of our hPSMC results to the inclusion or exclusion of repeat regions in the pseudodiploid genome. We compared the hPSMC results when including and removing repeat regions for three independent species pairs of varying phylogenetic distance. These included a shallow divergence (bottlenose and Indo-Pacific bottlenose dolphins), medium divergence (beluga and narwhal), and deep divergence (bottlenose dolphin and beluga) (Supplementary Figs. S5 - S7). For all species pairs, results showed that pre-divergence Ne is almost identical, and the exponential increase in Ne is just slightly more recent when removing repeat regions, compared to when repeat regions are included. This gives us confidence that the inclusion of repeats did not greatly alter our results.

To add independent evidence for continued lineage sorting/gene flow for an extended period after initial divergence, we compared relative divergence time between killer whale, Pacific white-sided dolphin, and long-finned pilot whale based on the species tree and a set of alternative topologies (Supplementary Fig. S8). We focused on Delphinidae, due to the large number of loci per alternative topology (Supplementary Tables S1, S2, S3, and S4). By assuming ILS and gene flow are the dominant forces behind gene-tree discordance, we can uncover information about the timing of ILS and gene flow events among lineages, by isolating the loci that produce each topology (Mendes and Hahn, 2016). In agreement with our hPSMC results, this analysis showed that ILS/gene flow continued for a long time after initial divergence. For example, we observed that the killer whale diverged from all other Delphinidae at a relative divergence time of 0.45 (45% of the divergence time of Delphinoidea and the baiji) in the consensus topology (Supplementary Fig. S8A). In an alternative topology, the killer whale was placed as sister to the Pacific white-sided dolphin (Supplementary Fig. S8B); despite still diverging from the remaining Delphinidae at approximately the same relative timing (0.42), it diverged from the Pacific white-sided dolphin at a relative divergence time of 0.25. As we assumed the alternative topologies only arose due to ILS and/or gene flow, this suggested lineage sorting and/or gene flow continued along ^~^40% of the post-divergence branch length. This estimate was qualitatively equivalent to that made using hPSMC (minimally 43%). Similarly, long periods of post-divergence lineage sorting/gene flow were observed when investigating topologies with the killer whale and long-finned pilot whale as sister species (Supplementary Fig. S8C, ^~^43%), and with the Pacific white-sided dolphin and long-finned pilot whale as sister species (Supplementary Fig. S8D, ^~^37%). As the results here included alternative topologies that likely arose due to both ILS and gene flow, we propose that the numbers present a more conservative estimate. One would expect ILS to be a more prevalent force behind discordances shortly after the species’ divergence, whereas gene flow can occur after many generations. Therefore, if we could more confidently disentangle alternative topologies arising due to ILS from those arising due to gene flow, we would expect much more recent relative divergence times for loci that underwent gene flow.

In summary, by combining findings from several analyses, and with the knowledge that interspecific hybridisation is still ongoing between many of the lineages studied here, we suggest that both ILS and gene flow played a major role over extended periods of time, in the speciation of Delphinoidea.

### Interspecific hybridisation

Making inferences as to what biological factors lead to interspecific hybridisation is challenging, as many variables may play a role. One hypothesis is that interspecific hybridization may occur at a higher rate during periods of low abundance, when a given species encounters only a limited number of conspecifics (Edwards *et al*., 2011; Crossman, Taylor and Barrett-Lennard, 2016; Westbury, Petersen and Lorenzen, 2019). When considering species that have not yet undergone sufficient divergence preventing their ability to hybridise, individuals may mate with a related species, instead of investing energy in finding a relatively rarer conspecific mate.

To explore the relationship between susceptibility to interspecific hybridisation and population size, we calculated the level of genome-wide genetic diversity for each species, as a proxy for their Ne (Fig. 4A). Narwhal, killer whale, beluga, and long-finned pilot whale had the lowest diversity levels, respectively, and should therefore be more susceptible to interspecific hybridization events. A beluga/narwhal hybrid has been reported (Skovrind *et al*., 2019), as has hybridisation between long-finned and short-finned pilot whales (Miralles *et al*., 2016). However, hybrids between species with high genetic diversity, including harbour porpoise (Willis *et al*., 2004), Indo-Pacific bottlenose dolphin (Baird *et al*., 2012), and bottlenose dolphin (Herzingl and Johnsonz, 1997; Espada *et al*., 2019), have also been reported, suggesting genetic diversity alone is not a good proxy for susceptibility to hybridisation.

**Figure 4:**
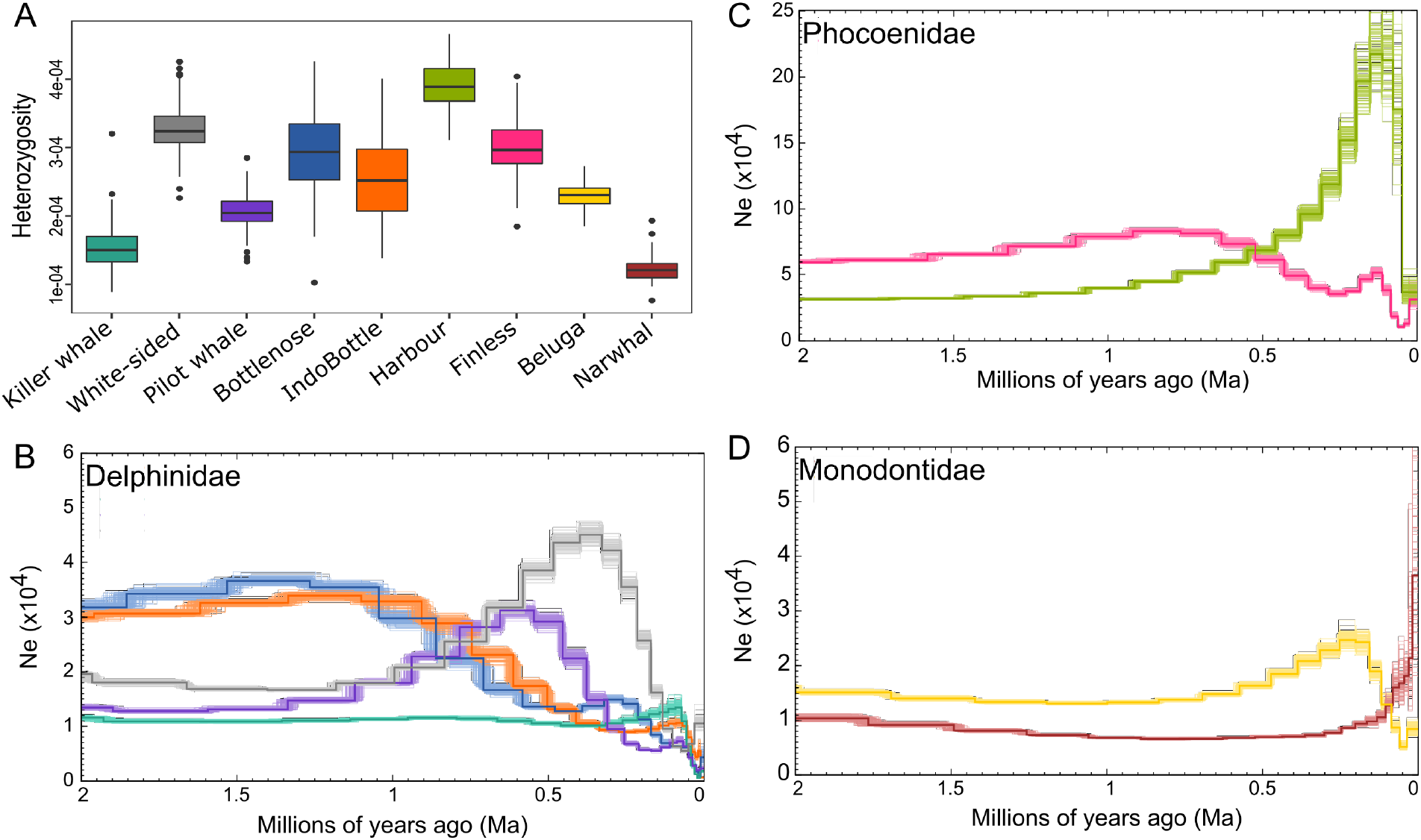
Autosome-wide heterozygosity and demographic histories over the past two million years. (A) Autosome-wide levels of heterozygosity calculated in 20 Mb sliding windows. (B-D) Demographic history of all studied species within (B) Delphinidae, (C) Phocoenidae, and (D) Monodontidae, estimated using PSMC. Thick coloured lines show estimated demographic trajectory, faded lines show bootstrap support values. Colours of B-D correspond to species’ colour from A.

To investigate the effect of interspecific gene flow on Ne, we estimated changes in intraspecific genetic diversity through time (Fig. 4B-D). The modelled demographic trajectories, using a Pairwise Sequentially Markovian Coalescent model (PSMC), span the past two million years. We could therefore assess the relationship for the three species pairs, where the putative interval for the cessation of lineage sorting/gene flow was contained within this period: harbour/finless porpoise (Phocoenidae), beluga/narwhal (Monodontidae), and bottlenose/Indo-Pacific bottlenose dolphin (Delphinidae) (Fig. 3).

In the harbour porpoise, we observed an increase in Ne beginning ^~^1 Ma, the rate of which increased further ^~^0.5 Ma (Fig. 4C). We observed a similar pattern in belugas; an increase in Ne ^~^1 Ma, relatively soon after the proposed cessation of gene flow with narwhals ^~^1.8 - 1.2 Ma (Fig. 4D). Although Ne may reflect abundance, it is also influenced by several other factors, including population connectivity and gene flow. If gene flow explained our changes in Ne, we would therefore expect a decrease in Ne after gene flow ceased, but instead we observed an increase. An increase in Ne may coincide with an increase in relative abundance, which would increase the number of potential conspecific mates, and in turn reduce the level of interspecific gene flow. However, this is difficult to say for certain without more information on abundances through time.

We observed a different pattern in the bottlenose/Indo-Pacific bottlenose dolphins. We found a relatively high population size during the period of gene flow in both species; Ne declines ^~^1 - 0.5 Ma, coinciding with the putative end of gene flow ^~^1.2 - 0.4 Ma. The decline in Ne could either reflect a decline in abundance, or a loss of connectivity between the two species. In the latter, we expect levels of intraspecific diversity (and thereby inferred Ne) to decline with the cessation of gene flow, even if absolute abundances did not change. This is indeed suggested by our data, which showed both species undergoing the decline simultaneously, indicative of a common cause.

Seven of the nine Delphinoidea genomes investigated showed a similar pattern of a rapid decline in Ne starting ^~^150-100 thousands of years ago (kya) (Fig. 4B-D; the exceptions are Pacific white-sided dolphin and narwhal). This concurrent decline could represent actual population declines across species, or, alternatively, simultaneous reductions in connectivity among populations within each species. Based on similar PSMC analyses, a decline in Ne at this time has also been reported in four baleen whale species (Árnason *et al*., 2018). Therefore, the species-wide pattern may reflect climate-driven environmental change. The period of 150-100 kya overlaps with the onset of the last interglacial, when sea levels increased to levels as high, if not higher, than at present (Polyak *et al*., 2018), and which may have had a marine-wide effect on both population connectivity and sizes. The unique life histories, distribution, and ecology of the cetacean species suggests that a combination of both decreased population connectivity and population sizes across the different studied species. A similar marine-wide effect has been observed among baleen whales and their prey species in the Southern and North Atlantic Oceans during the Pleistocene-Holocene climate transition (12-7 kya) (Cabrera *et al*., 2022). These results indicate that past marine-wide environmental shifts have driven demographic changes in population across multiple marine species.

Although speculative, we suggest that recent species-wide declines associated with the onset of the last glacial period, may have facilitated the resurgence of hybridization between some of the nine Delphinoidea species analysed. If interspecific hybridisation has increased after these declines, species may already be sufficiently differentiated that offspring fertility is reduced. Even if offspring are fertile, the high level of differentiation between species may mean hybrids are unable to occupy either parental niche (Skovrind *et al*., 2019) and are strongly selected against. A lack of significant contribution from recent hybrids to the parental gene pools may be why we observe contemporary hybrids, but do not find evidence of this in our analyses.

## Conclusions

Allopatric speciation is generally considered the most common mode of speciation, as the absence of gene flow due to geographic isolation can most easily explain the evolution of ecological, behavioural, morphological, or genetic differences between populations (Norris and Hull, 2012). However, our findings suggest that within Delphinoidea, speciation in the presence of gene flow was commonplace, consistent with sympatric/parapatric speciation, or allopatric speciation and secondary contact.

The ability for gene flow events to occur long after initial divergence may also explain the presence of contemporaneous hybrids between several species. In parapatric speciation, genetic isolation is achieved relatively early due to geographical and biological isolation, but species develop complete reproductive isolation relatively slowly, through low levels of migration or secondary contact events (Norris and Hull, 2012). The prevalence of this mode of speciation in cetaceans, as suggested by our study and previous genomic analyses (Árnason *et al*., 2018; Moura *et al*., 2020), may reflect the low energetic costs of dispersing across large distances in the marine realm (Williams, 1999; Fish, Howle and Murray, 2008) and the relative absence of geographic barriers preventing such dispersal events (Palumbi, 1994). Both factors are believed to be important in facilitating long-distance (including inter-hemispheric and inter-oceanic) movements in many cetacean species (Stone, Florez-Gonzalez and Katona, 1990).

Our study shows that speciation in Delphinoidea was a complex process and involved multiple ecological and evolutionary factors. Our results take a step towards resolving the enormous complexity of speciation within this superfamily, through a multifaceted analysis of nuclear genomes. Our study underscores the challenges of accurately interpreting some results, potentially due to the high levels of divergence between the target species and amplified by rapid diversification where ILS is likely pervasive, and where introgression among ancestral lineage was also likely. Moreover, while we make inferences based on a genome-wide dataset, certain regions of the genome may have a greater contribution to reproductive isolation than others, e.g. sex chromosomes and regions of reduced recombination (Payseur and Rieseberg, 2016). By using the hypotheses we form about general patterns and major processes of gene flow and speciation uncovered in our data, we hope that future studies may be able to build on our results to make more specific inferences as to the genomics of speciation in Delphinoidea, as additional genomic data and new methodologies for data analysis become available.

## Materials and Methods

### Data collection

We downloaded the assembled genomes and raw sequencing reads from nine toothed whales from the superfamily Delphinoidea. The data included five Delphinidae: Pacific white-sided dolphin (NCBI Biosample: SAMN09386610), Indo-Pacific bottlenose dolphin (NCBI Biosample: SAMN06289676), bottlenose dolphin (NCBI Biosample: SAMN09426418), killer whale (NCBI Biosample: SAMN01180276), and long-finned pilot whale (NCBI Biosample: SAMN11083132); two Phocoenidae: harbour porpoise (Autenrieth *et al*., 2018) and finless porpoise (NCBI Biosample: SAMN02192673); and two Monodontidae: beluga (NCBI Biosample: SAMN06216270) and narwhal (NCBI Biosample: SAMN10519625). To avoid reference biases, where reads more similar to the reference map more successfully than more divergent reads, artificially inflating signals of genetic similarities between a highly divergent outgroup and an ingroup species used as mapping reference (Liu *et al*., 2021), we downloaded the assembled outgroup baiji genome (Genbank accession code: GCF_000442215.1) as mapping reference in the gene flow analyses. Delphinoidea and the baiji diverged ^~^24.6 Ma (95% CI 25.2 - 23.8 Ma) (McGowen *et al*., 2020).

### Initial data filtering

To determine which scaffolds were most likely autosomal in origin, we identified putative sex chromosome scaffolds for each genome through synteny, and omitted them from further analysis. We found putative sex chromosome scaffolds in all ten assemblies by aligning them to the Cow X (Genbank accession: CM008168.2) and Human Y (Genbank accession: NC_000024.10) chromosomes. Alignments were performed using satsuma synteny v2.1 (Grabherr *et al*., 2010) with default parameters. Since short scaffolds have a higher likelihood of including assembly errors, we also removed scaffolds smaller than 100 kb from all downstream analyses.

### Mapping

We trimmed adapter sequences from all raw reads using skewer v0.2.2 (Jiang *et al*., 2014). We mapped the trimmed reads to the baiji for downstream gene flow analyses, and to the species-specific reference genome for downstream demographic history and genetic diversity analyses using BWA v0.7.15 (Li and Durbin, 2009) and the mem algorithm. We parsed the output and removed duplicates and reads with a mapping quality lower than 30 with SAMtools v1.6 (Li *et al*., 2009). Mapping statistics can be found in supplementary tables S8 and S9.

### Sliding-window phylogeny

For the sliding-window phylogenetic analysis, we created fasta files for all individuals mapped to the baiji genome using a consensus base call (-dofasta 2) approach in ANGSD v0.921 (Korneliussen, Albrechtsen and Nielsen, 2014), and specifying the following filters: minimum read depth of 5 (-mininddepth 5), minimum mapping quality of 30 (-minmapq 30), minimum base quality (-minq 30), only consider reads that map to one location uniquely (-uniqueonly 1), and only include reads where both mates map (-only_proper_pairs 1). All resultant fasta files, together with the assembled baiji genome, were aligned, and sites where any individual had more than 50% missing data were filtered before performing maximum likelihood phylogenetic analyses in a non-overlapping sliding-window approach using RAxML v8.2.10 (Stamatakis, 2014). We performed this analysis four times independently, specifying a different window size each time (50 kb, 100 kb, 500 kb, and 1 Mb). We used RAxML with default parameters and a GTR+G substitution model. Using the trees from each window, we estimated the species tree under the multi-species coalescent using ASTRAL-III (Zhang *et al*., 2018), and extracted the proportion of gene trees supporting each branch using PHYLIP (Felsenstein, 2005). We also visualised all trees of equal sized windows using DensiTree (Bouckaert, 2010).

We tested whether discordant phylogenetic topologies may be linked to GC content in the 50kb windows. To do this, we calculated the GC content for each window and binned the windows into three bins: The 33% with the lowest levels of GC content, the 33% with intermediate levels, and the 33% with the highest levels of GC content.

### Quantifying Introgression via Branch Lengths (QuIBL)

To test hypotheses of whether phylogenetic discordance between all possible triplets can be explained by incomplete lineage sorting (ILS) alone, or by a combination of ILS and gene flow, we implemented QuIBL (Edelman *et al*., 2019) in two different datasets. The first dataset leveraged the results of the above 50 kb-window analysis, by taking every twentieth tree from the 50kb sliding-window analysis and running it through QuIBL. The second dataset was created specifically for this test, and contained topologies generated from 20 kb windows with a 1 Mb slide using the phylogenetic methods mentioned above. We ran QuIBL specifying the baiji as the overall outgroup (totaloutgroup), to test either ILS or ILS with gene flow (numdistributions 2), the number of total EM steps as 50 (numsteps), and a likelihood threshold of 0.01. We determined the significance of gene flow by comparing the BIC1 (ILS alone) and BIC2 (ILS and gene flow). When BIC2 was lower than BIC1, with a difference of > 10, we assumed incongruent topologies arose due to both ILS and gene flow. Triplet topologies supporting the species tree, and those that had < 5 alternative topologies, were excluded from interpretations.

### D-statistics

To test for signs of gene flow in the face of ILS, we ran D-statistics (Green *et al*., 2010; Durand *et al*., 2011) using all individuals mapped to the baiji genome in ANGSD, and using a consensus base call approach (- doabbababa 2), specifying the baiji sequence as the ancestral outgroup sequence, and the same filtering as for the fasta file construction with the addition of setting the block size as 1Mb (-blocksize). Significance of the results was evaluated using a block jackknife approach with the Rscript provided in the ANGSD package. |Z| > 3 was deemed significant.

### D-foil

As D-statistics only tests for the presence and not the direction of gene flow, we ran D-foil (Pease and Hahn, 2015), an extended version of the D-statistic, which is a five-taxon test for gene flow, making use of all four combinations of the potential D-statistics topologies. For this analysis, we used the same fasta files constructed above, which we converted into an mvf file using MVFtools (Pease and Rosenzweig, 2018). We specified the 5-taxa [[H1, H2], [H3, H4], baiji], for all possible combinations, following the species tree (Fig. 1) and a 100 kb window size. All scaffolds were trimmed to the nearest 100 kb to avoid the inclusion of windows shorter than 100 kb. The significance of each window was separately assessed by a chi-squared goodness-of-fit test within the software.

### The *f*-branch statistic

To aid in the interpretation of the multitude of D-statistics comparisons, we implemented the *f*-branch test (Malinsky *et al*., 2018; Malinsky, Matschiner and Svardal, 2021) to uncover correlations between results that may indicate ancestral gene flow events. For this analysis, we needed a variant call file (VCF). However, the raw sequencing reads for the baiji are not available. To overcome this, we simulated 100 million 150 bp reads from the assembled genome using SAMtools wgsim, which we mapped back to the baiji assembly using the same mapping parameters specified above. We constructed a multi-individual VCF of all individuals mapped to the baiji using bcftools mpileup, and filtered said VCF file to only include SNPs using BCFtools call and the - mv parameter, resulting in 138,715,767 sites for downstream analyses. We ran the multi-individual VCF through Dtrios in Dsuite v0.4 r43 (Malinsky, Matschiner and Svardal, 2021) and specified the species tree as the most common topology from our sliding window analyses, and otherwise default parameters. We ran the output from Dtrios through *f*-branch and visualised the output using the dtools.py script from Dsuite. To assess whether sex chromosomes may support a different scenario of gene flow events, we also ran the *f*-branch on scaffolds >1 Mb aligning to the X chromosome which gave us 3,728,572 sites. Although the default parameters for the *f*-branch statistic in Dsuite only consider *f*b with p<0.01, we also assessed statistical significance of *f*b using a block Jack-knife approach by including the -Z parameter when running the *f*-branch statistic in Dsuite. A Z score |Z|>3 was considered as significant.

### Mutation rate estimation

For use in the downstream demographic analyses, we computed the mutation rate per generation for each species. To do this, we estimated the pairwise distances between all ingroup species mapped to the baiji, using a consensus base call in ANGSD (-doIBS 2), and applying the same filters as above, with the addition of only considering sites in which all individuals were covered (-minInd). The pairwise distances used in this calculation were those from the closest lineage to the species of interest (Supplementary Tables S10 and S11). The mutation rates per generation were calculated using the resultant pairwise distance as follows: mutation rate = pairwise distance x generation time / 2 x divergence time. Divergence times were taken from the full dataset 10-partition auto-correlated rate (mean) values from McGowen *et al*. (2020) (Supplementary Table S11). Generation times were taken from previously published data (Supplementary Table S12).

### Cessation of lineage sorting and/or gene flow

To estimate when lineage sorting and/or gene flow may have ceased between each species pair, we used the F1-hybrid PSMC (hPSMC) approach (Cahill *et al*., 2016). As input we used the haploid consensus sequences mapped to the baiji that were created for the phylogenetic analyses. Despite the possibility of producing consensus sequences when mapping to conspecific reference genomes, we chose the baiji for all comparisons, as previous analyses have shown the choice of reference genome does not influence hPSMC results (Westbury, Petersen and Lorenzen, 2019; Moodley *et al*., 2020). We merged the haploid sequences from each possible species pair into pseudo-diploid sequences using the scripts available in the hPSMC toolsuite. We independently ran each resultant species pair pseudo-diploid sequences through PSMC, specifying atomic intervals 4+25*2+4+6. We plotted the results using the average (i) mutation rate per generation and (ii) generation time for each species pair being tested. From the output of this analysis, we visually estimated the pre-divergence Ne of each hPSMC plot (i.e. Ne prior to the point of asymptotic increase in Ne) to be used as input for downstream simulations. Based on these empirical results, we ran simulations in ms (Hudson, 2002) using the estimated pre-divergence Ne, and various predefined divergence times, to find the interval in which gene flow may have ceased between a given species pair. The time intervals and pre-divergence Ne for each species pair used for the simulations can be seen in supplementary table S7. The ms commands were produced using the scripts available in the hPSMC toolsuite. We plotted the simulated and empirical hPSMC results to find the simulations with an asymptotic increase in Ne closest to, but not overlapping with, the empirical data. The predefined divergence times of the simulations showing this pattern within 1.5x and 10x of the pre-divergence Ne were taken as the time interval in which gene flow ceased.

We repeated the above analysis for three species pairs: bottlenose/Indo-Pacific bottlenose dolphins, beluga/narwhal, and beluga/bottlenose dolphin, but with an additional step, where we masked repeat elements of the haploid genomes using bedtools v2.26.0 (Quinlan, 2014) and the repeat annotations available on Genbank. Once we masked the repeat elements, we re-ran the hPSMC analysis as above.

### Relative divergence times in Delphinidae

To further examine the timing of the ending of lineage sorting and/or gene flow, we performed phylogenetic inferences to uncover the relative divergence times on subsets of genomic loci showing alternative topologies in Delphinidae. To do this, we masked repeats in the same fasta files used for our other phylogenetic analyses using the baiji Genbank annotation and bedtools (Quinlan, 2014). We extracted 1 kb windows with a 1 Mb slide from the aligned fasta files and only kept loci containing less than 50% missing data for any individual. We separated our data set into the loci that supported each of four sets of relationships. These included loci that supported (i) the consensus species tree (n = 109), (ii) the Pacific white-sided dolphin as sister to the killer-whale (n = 84), (iii) the Pacific white-sided dolphin as sister to the clade of bottlenose dolphins, with the long-finned pilot and killer whales in a monophyletic clade as sisters to this group (n = 48), and (iv) the Pacific white-sided dolphin as sister to the long-finned pilot whale (n = 59).

As focal species, we selected to test the Pacific white-sided dolphin, killer whale, and long-finned pilot whale, as they showed the highest number of discordances, allowing for a more balanced comparison of divergence-time estimates among different topologies. For each of the four sets of loci, we inferred the relative divergence times across our samples of Delphinidae, also including the beluga and the baiji in the taxon set. We analysed each data set independently, constrained the tree topology to that of the corresponding set of loci, and constrained the age of the root to 1. We performed Bayesian dating using a GTR+Γ substitution model and an uncorrelated-gamma relaxed clock model in MCMCtree, as implemented in PAML v4.8 (Yang, 2007). The posterior distribution was approximated using Markov chain Monte Carlo (MCMC) sampling, with samples drawn every 10^3^ MCMC steps over 10^7^ steps, after discarding a burn-in phase of 10^5^ steps. Convergence to the stationary distribution was verified by comparing parameter estimates from two independent analyses, and confirming that effective sample sizes were above 200 for all sampled parameters.

### Heterozygosity

As a proxy for species-level genetic diversity, we estimated autosome-wide heterozygosity for each of the nine Delphinoidea species. We estimated autosomal heterozygosity using allele frequencies (-doSaf 1) in ANGSD (Korneliussen, Albrechtsen and Nielsen, 2014), taking genotype likelihoods into account (-GL 2) and specifying the same filters as for the fasta file construction, with the addition of adjusting quality scores around indels (-baq 1). To ensure comparability between genomes of differing coverage, we uniquely set the subsample filter (-downSample) for each individual to result in a 20x genome-wide coverage. Heterozygosity was computed from the output of this using realSFS from the ANGSD toolsuite and specifying 20 Mb windows of covered sites (-nSites).

### Demographic reconstruction

To determine the demographic histories of all nine species over a two million year time scale, we ran a Pairwise Sequentially Markovian Coalescent model (PSMC) (Li and Durbin, 2011) on each diploid genome independently. We called diploid genome sequences using SAMtools and BCFtools v1.6 (Narasimhan *et al*., 2016), specifying a minimum quality score of 20 and minimum coverage of 10. We ran PSMC specifying atomic intervals 4+25*2+4+6 and performed 100 bootstrap replicates to investigate support for the resultant demographic trajectories. PSMC outputs were plotted using species-specific mutation rates and generation times (Supplementary Table S12).

## Acknowledgements

We would like to thank all those contributing to the ever-increasing abundance of publicly available genomic resources. Without the availability of such data, our study would not have been possible. We would also like to thank Michael Fontaine, Christelle Fraïsse, Camille Roux, Andrew Foote, and Simon Martin for their helpful input to previous versions of this manuscript. Version 7 of this preprint has been peer-reviewed and recommended by Peer Community In Evolutionary Biology (https://doi.org/10.24072/pci.evolbiol.100139).

## Data, script and code availability

All data for this manuscript were downloaded from Genbank with accession codes provided in the material and methods. Previously unpublished scripts and example commands for the analyses performed in this manuscript can be found at https://doi.org/10.5281/zenodo.6580299.

## Supplementary information availability

Supplementary materials are available online: https://doi.org/10.5281/zenodo.6581045. This includes supplementary figures S1 - S8, supplementary tables S1 - S12, and supplementary results - hPSMC.

## Author contributions

Conceptualization, MVW; Formal analysis, MVW, AAC, AR-I, BDC, DAD, SH; Writing – Original Draft MVW; Writing – Review & Editing, All authors; Supervision, MVW, EDL; Funding Acquisition, EDL

## References

Árnason, Ú. et al. (2018) ‘Whole-genome sequencing of the blue whale and other rorquals finds signatures for introgressive gene flow’, Science advances, 4(4), p. eaap9873. http://doi.org/10.1126/sciadv.aap9873

Autenrieth, M. et al. (2018) ‘High-quality whole-genome sequence of an abundant Holarctic odontocete, the harbour porpoise (Phocoena phocoena)’, Molecular ecology resources, 18(6), pp. 1469–1481. http://doi.org/10.1111/1755-0998.12932

Baird, R.W. et al. (2012) ‘Population structure of island-associated dolphins: Evidence from mitochondrial and microsatellite markers for common bottlenose dolphins (Tursiops truncatus) in the main Hawaiian Islands’, Marine Mammal Science, 25(2), pp. 251–274. http://doi.org/10.1111/j.1748-7692.2008.00257.x

Barlow, A. et al. (2018) ‘Partial genomic survival of cave bears in living brown bears’, Nature ecology & evolution, 2(10), pp. 1563–1570. http://doi.org/10.1038/s41559-018-0654-8

Bierne, N., Bonhomme, F. and David, P. (2003) ‘Habitat preference and the marine-speciation paradox’, Proceedings of the Royal Society B: Biological Sciences, 270(1522), pp. 1399–1406. http://doi.org/10.1098/rspb.2003.2404

Bouckaert, R.R. (2010) ‘DensiTree: making sense of sets of phylogenetic trees’, Bioinformatics, 26(10), pp. 1372–1373. http://doi.org/10.1093/bioinformatics/btq110

Butlin, R.K. and Smadja, C.M. (2018) ‘Coupling, Reinforcement, and Speciation’, The American naturalist, 191(2), pp. 155–172. http://doi.org/10.1086/695136

Cabrera, A.A. et al. (2022) ‘Strong and lasting impacts of past global warming on baleen whales and their prey’, Global change biology, 28(8), pp. 2657–2677. http://doi.org/10.1111/gcb.16085

Cahill, J.A. et al. (2016) ‘Inferring species divergence times using pairwise sequential Markovian coalescent modelling and low-coverage genomic data’, Proceedings of the Royal Society B: Biological Sciences, 371(1699). http://doi.org/10.1098/rstb.2015.0138

Campbell, C.R. and Poelstra, J.W. (2018) ‘What is Speciation Genomics? The roles of ecology, gene flow, and genomic architecture in the formation of species’, Biological journal of the Linnean Society. Linnean Society of London, 124(4), pp. 561–583. http://doi.org/10.1093/biolinnean/bly063

Crossman, C.A., Taylor, E.B. and Barrett-Lennard, L.G. (2016) ‘Hybridization in the Cetacea: widespread occurrence and associated morphological, behavioral, and ecological factors’, Ecology and evolution, 6(5), pp. 1293–1303. http://doi.org/10.1002/ece3.1913

Durand, E.Y. et al. (2011) ‘Testing for ancient admixture between closely related populations’, Molecular biology and evolution, 28(8), pp. 2239–2252. http://doi.org/10.1093/molbev/msr048

Edelman, N.B. et al. (2019) ‘Genomic architecture and introgression shape a butterfly radiation’, Science, 366(6465), pp. 594–599. http://doi.org/10.1126/science.aaw2090

Edwards, C.J. et al. (2011) ‘Ancient hybridization and an Irish origin for the modern polar bear matriline’, Current biology: CB, 21(15), pp. 1251–1258. http://doi.org/10.1016/j.cub.2011.05.058

Espada, R. et al. (2019) ‘Hybridization in the wild between Tursiops truncatus (Montagu 1821) and Delphinus delphis (Linnaeus 1758)’, PloS one, 14(4), p. e0215020. http://doi.org/10.1371/journal.pone.0215020

Feder, J.L., Egan, S.P. and Nosil, P. (2012) ‘The genomics of speciation-with-gene-flow’, Trends in genetics: TIG, 28(7), pp. 342–350. http://doi.org/10.1016/j.tig.2012.03.009

Felsenstein, J. (2005) ‘PHYLIP (Phylogeny Inference Package) version 3.6’. Department of Genome Sciences, University of Washington, Seattle. Available at: http://evolution.genetics.washington.edu/phylip/

Fish, F.E., Howle, L.E. and Murray, M.M. (2008) ‘Hydrodynamic flow control in marine mammals’, Integrative and comparative biology, 48(6), pp. 788–800. http://doi.org/10.1093/icb/icn029

Foote, A.D. et al. (2011) ‘Out of the Pacific and back again: insights into the matrilineal history of Pacific killer whale ecotypes’, PLoS One, 6(9), e24980. http://doi.org/10.1371/journal.pone.0024980

Foote, A.D. (2018) ‘Sympatric Speciation in the Genomic Era’, Trends in ecology & evolution, 33(2), pp. 85–95. http://doi.org/10.1016/j.tree.2017.11.003

Foote, A.D. and Morin, P.A. (2015) ‘Sympatric speciation in killer whales?’, Heredity, 114(6), pp. 537–538. http://doi.org/10.1038/hdy.2014.120

Grabherr, M.G. et al. (2010) ‘Genome-wide synteny through highly sensitive sequence alignment: Satsuma’, Bioinformatics, 26(9), pp. 1145–1151. http://doi.org/10.1093/bioinformatics/btq102

Green, R.E. et al. (2010) ‘A draft sequence of the Neandertal genome’, Science, 328(5979), pp. 710–722. http://doi.org/10.1126/science.1188021

Gridley, T. et al. (2018) ‘Hybridization in bottlenose dolphins—A case study of Tursiops aduncus × T. truncatus hybrids and successful backcross hybridization events’, PloS one, 13(9), p. e0201722. http://doi.org/10.1371/journal.pone.0201722

Herzingl, D.L. and Johnsonz, C.M. (1997) ‘Interspecific interactions between Atlantic spotted dolphins (Stenella frontalis) and bottlenose dolphins (Tursiops truncatus) in the Bahamas 1985-1995’, Aquatic Mammals, 23, pp. 85–99. Available at: https://www.aquaticmammalsjournal.org/share/AquaticMammalsIssueArchives/1997/AquaticMammals_23-02/23-02_Herzing.pdf

Hudson, R.R. (2002) ‘Generating samples under a Wright–Fisher neutral model of genetic variation’, Bioinformatics, 18(2), pp. 337–338. http://doi.org/10.1093/bioinformatics/18.2.337

Jiang, H. et al. (2014) ‘Skewer: a fast and accurate adapter trimmer for next-generation sequencing paired-end reads’, BMC bioinformatics, 15, p. 182. http://doi.org/10.1186/1471-2105-15-182

Korneliussen, T.S., Albrechtsen, A. and Nielsen, R. (2014) ‘ANGSD: Analysis of Next Generation Sequencing Data’, BMC bioinformatics, 15, p. 356. http://doi.org/10.1186/s12859-014-0356-4

Lartillot, N. (2013) ‘Phylogenetic patterns of GC-biased gene conversion in placental mammals and the evolutionary dynamics of recombination landscapes’, Molecular biology and evolution, 30(3), pp. 489–502. http://doi.org/10.1093/molbev/mss239

Leaché, A.D. et al. (2014) ‘The influence of gene flow on species tree estimation: a simulation study’, Systematic biology, 63(1), pp. 17–30. http://doi.org/10.1093/sysbio/syt049

Li, H. et al. (2009) ‘The Sequence Alignment/Map format and SAMtools’, Bioinformatics, 25(16), pp. 2078–2079. http://doi.org/10.1093/bioinformatics/btp352

Li, H. and Durbin, R. (2009) ‘Fast and accurate short read alignment with Burrows–Wheeler transform’, Bioinformatics, 25(14), pp. 1754–1760. http://doi.org/10.1093/bioinformatics/btp324

Li, H. and Durbin, R. (2011) ‘Inference of human population history from individual whole-genome sequences’, Nature, 475(7357), pp. 493–496. http://doi.org/10.1038/nature10231

Liu, S. et al. (2014) ‘Population genomics reveal recent speciation and rapid evolutionary adaptation in polar bears’, Cell, 157(4), pp. 785–794. http://doi.org/10.1016/j.cell.2014.03.054

Liu, S. et al. (2021) ‘Ancient and modern genomes unravel the evolutionary history of the rhinoceros family’, Cell, 184(19), pp. 4874–4885.e16. http://doi.org/10.1016/j.cell.2021.07.032

Malinsky, M. et al. (2018) ‘Whole-genome sequences of Malawi cichlids reveal multiple radiations interconnected by gene flow’, Nature ecology & evolution, 2(12), pp. 1940–1955. http://doi.org/10.1038/s41559-018-0717-x

Malinsky, M., Matschiner, M. and Svardal, H. (2021) ‘Dsuite - Fast D-statistics and related admixture evidence from VCF files’, Molecular ecology resources, 21(2), pp. 584–595. http://doi.org/10.1111/1755-0998.13265

Martin, S.H., Davey, J.W. and Jiggins, C.D. (2015) ‘Evaluating the use of ABBA–BABA statistics to locate introgressed loci’, Molecular biology and Evolution, 32(1), pp. 244–257. http://doi.org/10.1093/molbev/msu269

McGowen, M.R. et al. (2020) ‘Phylogenomic Resolution of the Cetacean Tree of Life Using Target Sequence Capture’, Systematic biology, 69(3), pp. 479–501. http://doi.org/10.1093/sysbio/syz068

Mendes, F.K. and Hahn, M.W. (2016) ‘Gene Tree Discordance Causes Apparent Substitution Rate Variation’, Systematic biology, 65(4), pp. 711–721. http://doi.org/10.1093/sysbio/syw018

Miralles, L. et al. (2016) ‘Interspecific Hybridization in Pilot Whales and Asymmetric Genetic Introgression in Northern Globicephala melas under the Scenario of Global Warming’, PloS one, 11(8), p. e0160080. http://doi.org/10.1371/journal.pone.0160080

Miyazaki, N. et al. (1992) ‘Osteological study of a hybrid between Tursiops truncatus and Grampus griseus’, Bulletin of the National Museum of Nature and Science. Series B, Botany, 18, pp. 79–94

Moodley, Y. et al. (2020) ‘Interspecific gene flow and the evolution of specialisation in black and white rhinoceros’, Molecular biology and evolution 37(11),pp. 3105–3117. http://doi.org/10.1093/molbev/msaa148

Moura, A.E. et al. (2015) ‘Phylogenomics of the killer whale indicates ecotype divergence in sympatry’, Heredity, 114(1), pp. 48–55. http://doi.org/10.1038/hdy.2014.67

Moura, A.E. et al. (2020) ‘Phylogenomics of the genus Tursiops and closely related Delphininae reveals extensive reticulation among lineages and provides inference about eco-evolutionary drivers’, Molecular phylogenetics and evolution, 146, p. 106756. http://doi.org/10.1016/j.ympev.2020.106756

Narasimhan, V. et al. (2016) ‘BCFtools/RoH: a hidden Markov model approach for detecting autozygosity from next-generation sequencing data’, Bioinformatics, 32(11), pp. 1749–1751. http://doi.org/10.1093/bioinformatics/btw044

Norris, R.D. and Hull, P.M. (2012) ‘The temporal dimension of marine speciation’, Evolutionary ecology, 26(2), pp. 393–415. http://doi.org/10.1007/s10682-011-9488-4

Palumbi, S.R. (1994) ‘Genetic divergence, reproductive isolation, and marine speciation’, Annual review of ecology and systematics, 25(1), pp. 547–572. http://doi.org/10.1146/annurev.es.25.110194.002555

Pamilo, P. and Nei, M. (1988) ‘Relationships between gene trees and species trees’, Molecular biology and evolution, 5(5), pp. 568–583. http://doi.org/10.1093/oxfordjournals.molbev.a040517

Payseur, B.A. and Rieseberg, L.H. (2016) ‘A genomic perspective on hybridization and speciation’, Molecular ecology, 25(11), pp. 2337–2360. http://doi.org/10.1111/mec.13557

Pease, J.B. and Hahn, M.W. (2015) ‘Detection and Polarization of Introgression in a Five-Taxon Phylogeny’, Systematic biology, 64(4), pp. 651–662. http://doi.org/10.1093/sysbio/syv023

Pease, J.B. and Rosenzweig, B.K. (2018) ‘Encoding Data Using Biological Principles: The Multisample Variant Format for Phylogenomics and Population Genomics’, IEEE/ACM transactions on computational biology and bioinformatics, 15(4), pp. 1231–1238. http://doi.org/10.1109/TCBB.2015.2509997

Polyak, V.J. et al. (2018) ‘A highly resolved record of relative sea level in the western Mediterranean Sea during the last interglacial period’, Nature geoscience, 11(11), pp. 860–864. http://doi.org/10.1038/s41561-018-0222-5

Quinlan, A.R. (2014) ‘BEDTools: The Swiss-Army Tool for Genome Feature Analysis’, Current protocols in bioinformatics, 47, pp. 11.12.1–34. http://doi.org/10.1002/0471250953.bi1112s47

Silva, J.M., Silva, F.J.L. and Sazima, I. (2005) ‘Two presumed interspecific hybrids in the genus Stenella (Delphinidae) in the Tropical West Atlantic’, Aquatic Mammals, 31(4), p. 468. http://doi.org/10.1578/AM.31.4.2005.468

Skovrind, M. et al. (2019) ‘Hybridization between two high Arctic cetaceans confirmed by genomic analysis’, Scientific reports, 9(1), p. 7729. http://doi.org/10.1038/s41598-019-44038-0

Slatkin, M. and Pollack, J.L. (2008) ‘Subdivision in an ancestral species creates asymmetry in gene trees’, Molecular biology and evolution, 25(10), pp. 2241–2246. http://doi.org/10.1093/molbev/msn172

Stamatakis, A. (2014) ‘RAxML version 8: a tool for phylogenetic analysis and post-analysis of large phylogenies’, Bioinformatics, 30(9), pp. 1312–1313. http://doi.org/10.1093/bioinformatics/btu033

Steeman, M.E. et al. (2009) ‘Radiation of extant cetaceans driven by restructuring of the oceans’, Systematic biology, 58(6), pp. 573–585. http://doi.org/10.1093/sysbio/syp060

Stone, G., Florez-Gonzalez, L. and Katona, S. (1990) ‘Whale migration record’, Nature, 346(6286), pp. 705–705. http://doi.org/10.1038/346705a0

Turelli, M., Barton, N.H. and Coyne, J.A. (2001) ‘Theory and speciation’, Trends in ecology & evolution, 16(7), pp. 330–343. http://doi.org/10.1016/s0169-5347(01)02177-2

Westbury, M.V. et al. (2020) ‘Hyena paleogenomes reveal a complex evolutionary history of cross-continental gene flow between spotted and cave hyena’, Science Advances, 6(11), p. eaay0456. http://doi.org/10.1126/sciadv.aay0456

Westbury, M.V., Petersen, B. and Lorenzen, E.D. (2019) ‘Genomic analyses reveal an absence of contemporary introgressive admixture between fin whales and blue whales, despite known hybrids’, PloS one, 14(9), p. e0222004. http://doi.org/10.1371/journal.pone.0222004

Williams, T.M. (1999) ‘The evolution of cost efficient swimming in marine mammals: limits to energetic optimization’, Philosophical Transactions of the Royal Society of London. Series B: Biological Sciences, 354(1380), pp. 193–201. http://doi.org/10.1098/rstb.1999.0371

Willis, P.M. et al. (2004) ‘Natural hybridization between Dall’s porpoises (Phocoenoides dalli) and harbour porpoises (Phocoena phocoena)’, Canadian journal of zoology, 82(5), pp. 828–834. http://doi.org/10.1139/z04-059

Yang, Z. (2007) ‘PAML 4: phylogenetic analysis by maximum likelihood’, Molecular biology and evolution, 24(8), pp. 1586–1591. http://doi.org/10.1093/molbev/msm088

Zhang, C. et al. (2018) ‘ASTRAL-III: polynomial time species tree reconstruction from partially resolved gene trees’, BMC bioinformatics, 19(Suppl 6), p. 153. http://doi.org/10.1186/s12859-018-2129-y

Zheng, Y. and Janke, A. (2018) ‘Gene flow analysis method, the D-statistic, is robust in a wide parameter space’, BMC bioinformatics, 19(1), p. 10. http://doi.org/10.1186/s12859-017-2002-4

