## Supplementary information for "A genomic assessment of the marine-speciation paradox within the toothed whale superfamily Delphinoidea" (from another version of this preprint: version 6, 2022-03-28)

**Supplementary table S1:** Proportions of the most frequent five topologies based on window sizes. NA - not in the five most frequent for that window size. Whitesided - Pacific white-sided dolphin, Pilotwhale - long-finned pilot whale, IndoBottlenose - Indo-Pacific bottlenose dolphin, Bottlenose - bottlenose dolphin, Killerwhale - killer whale, Beluga - beluga, Narwhal - narwhal, Harbour - harbour porpoise, Finless - finless porpoise, Baiji - Baiji (outgroup).

| 50kb | 100kb | 500kb | 1Mb | Topology |
|---|---|---|---|---|
| 0.24 | 0.32 | 0.64 | 0.79 | ((((Whitesided,(Pilotwhale,(IndoBottlenose,Bottlenose))),Killerwhale),((Beluga,Narwhal),(Harbour,Finless))),Baiji); |
| 0.14 | 0.14 | 0.09 | 0.05 | ((((Pilotwhale,(IndoBottlenose,Bottlenose)),(Whitesided,Killerwhale)),((Beluga,Narwhal),(Harbour,Finless))),Baiji); |
| 0.13 | 0.14 | 0.14 | 0.10 | ((((Pilotwhale,(Whitesided,(IndoBottlenose,Bottlenose))),Killerwhale),((Beluga,Narwhal),(Harbour,Finless))),Baiji); |
| 0.09 | 0.08 | 0.04 | 0.02 | (((((Pilotwhale,Whitesided),(IndoBottlenose,Bottlenose)),Killerwhale),((Beluga,Narwhal),(Harbour,Finless))),Baiji); |
| 0.08 | NA | NA | NA | ((((Killerwhale,(Pilotwhale,(IndoBottlenose,Bottlenose))), Whitesided),((Beluga,Narwhal),(Harbour,Finless))),Baiji); |
| NA | 0.07 | 0.03 | 0.02 | (((Whitesided,((Pilotwhale,(IndoBottlenose,Bottlenose)),Killerwhale)),((Beluga,Narwhal),(Harbour,Finless))),Baiji); |
| 0.69 | 0.76 | 0.94 | 0.98 | Top 5 topologies combined |

**Supplementary table S2:** Proportions of the most frequent five topologies based on GC content and a window size of 50kb. NA - not in the five most frequent for that window size. Whitesided - Pacific white-sided dolphin, Pilotwhale - long-finned pilot whale, IndoBottlenose - Indo-Pacific bottlenose dolphin, Bottlenose - bottlenose dolphin, Killerwhale - killer whale, Beluga - beluga, Narwhal - narwhal, Harbour - harbour porpoise, Finless - finless porpoise, Baiji - Baiji (outgroup).

| Low GC | Medium GC | High GC | Topology |
|---|---|---|---|
| 2814 | 3395 | 4227 | (((Killerwhale,(Whitesided,((IndoBottlenose,Bottlenose),Pilotwhale))),((Beluga,Narwhal),(Harbour,Finless))),Baiji); |
| 2023 | 2107 | 2085 | ((((Pilotwhale,(IndoBottlenose,Bottlenose)),(Whitesided,Killerwhale)),((Beluga,Narwhal),(Harbour,Finless))),Baiji); |
| 1740 | 1898 | 1976 | ((((Pilotwhale,(Whitesided,(IndoBottlenose,Bottlenose))),Killerwhale),((Beluga,Narwhal),(Harbour,Finless))),Baiji); |
| 1287 | 1289 | 1317 | (((((Pilotwhale,Whitesided),(IndoBottlenose,Bottlenose)),Killerwhale),((Beluga,Narwhal),(Harbour,Finless))),Baiji); |
| 1152 | NA | NA | ((((Whitesided,(IndoBottlenose,Bottlenose)),(Pilotwhale,Killerwhale)),((Beluga,Narwhal),(Harbour,Finless))),Baiji); |
| NA | 1190 | 1149 | (((Whitesided,((Pilotwhale,(IndoBottlenose,Bottlenose)),Killerwhale)),((Beluga,Narwhal),(Harbour,Finless))),Baiji); |

**Supplementary table S3:** QuIBL results when using every twentieth tree from the 50kb sliding window analysis - attached as spreadsheet. QuIBL analyses all triplet combinations ((A, B), C) in a given set of phylogenetic trees. Here we only present the alternative topologies within Delphinidae, that are in disagreement with the species tree, and may have arisen due to ILS or gene flow. The gene flow pair shows individuals A and B and outgroup is C. Two BIC scores are presented - one for ILS alone and one for ILS and gene flow. A BIC difference >10 suggests ILS and geneflow both as factors giving rise to the discordance topologies. % of total trees shows the percentage of all trees in the dataset having said triplet topology, whereas % of trees supporting topology explained by gene flow shows the percentage of the trees supporting said triplet topology that likely arose due to gene flow (based on branch length) instead of ILS. - attached as spreadsheet

**Supplementary table S4:** QuIBL results from trees constructed using 20kb windows with a 1Mb slide - attached as spreadsheet. QuIBL analyses all triplet combinations ((A, B), C) in a given set of phylogenetic trees. Here we only present the alternative topologies within Delphinidae, that are in disagreement with the species tree, and may have arisen due to ILS or gene flow. The gene flow pair shows individuals A and B and outgroup is C. Two BIC scores are presented - one for ILS alone and one for ILS and gene flow. A BIC difference >10 suggests ILS and geneflow both as factors giving rise to the discordance topologies. '% of total trees' shows the percentage of all trees in the dataset having said triplet topology. '% of trees supporting topology explained by gene flow' shows the percentage of the trees supporting said triplet topology that likely arose due to gene flow (based on branch length) instead of ILS. - attached as spreadsheet

**Supplementary table S5:** D-statistics results for all triplet combinations phylogenetically concurrent with our results shown in Figure 1. Baiji was used as the outgroup/ancestral sequence. A non-significant result ($|Z| < 3$) is indicated in bold. Colours indicate the family of the given individual. Red = Delphinidae, yellow = Phocoenidae, blue = Monodontidae.

| H1 | H2 | H3 | nABBA | nBABA | D-score | Z-score |
|---|---|---|---|---|---|---|
| Bottlenose | IndoBottlenose | Killer whale | 597,251 | 554,780 | 0.037 | 23.26 |
| Bottlenose | IndoBottlenose | Pilotwhale | 748,948 | 691,844 | 0.040 | 24.13 |
| Bottlenose | IndoBottlenose | Whitesided | 721,498 | 665,420 | 0.040 | 25.20 |
| Pilotwhale | Whitesided | Killer whale | 2,224,888 | 2,119,068 | 0.024 | 11.77 |
| Pilotwhale | Bottlenose | Killer whale | 1,998,297 | 1,795,444 | 0.053 | 26.15 |
| Pilotwhale | IndoBottlenose | Killer whale | 2,004,478 | 1,757,429 | 0.066 | 31.95 |
| Pilotwhale | Bottlenose | Whitesided | 2,490,189 | 2,051,579 | 0.097 | 42.67 |
| Pilotwhale | IndoBottlenose | Whitesided | 2,508,755 | 2,007,966 | 0.111 | 48.64 |
| Whitesided | Bottlenose | Killer whale | 2,111,742 | 2,014,525 | 0.024 | 11.88 |
| Whitesided | IndoBottlenose | Killer whale | 2,117,925 | 1,975,800 | 0.035 | 17.25 |
| Killer whale | Pilotwhale | Finless | 928,942 | 840,273 | 0.050 | 51.99 |

| | | | | | | |
|---|---|---|---|---|---|---|
| Killer whale | Whitesided | Finless | 924,323 | 829,525 | 0.054 | 56.12 |
| Killer whale | Pilotwhale | Harbour porpoise | 959,748 | 851,885 | 0.060 | 60.74 |
| Killer whale | Whitesided | Harbour porpoise | 956,686 | 840,318 | 0.065 | 65.46 |
| Killer whale | Bottlenose | Finless | 942,684 | 757,495 | 0.109 | 107.12 |
| Killer whale | Bottlenose | Harbour porpoise | 974,032 | 767,636 | 0.119 | 116.98 |
| Killer whale | IndoBottlenose | Finless | 943,526 | 728,185 | 0.129 | 120.99 |
| Killer whale | IndoBottlenose | Harbour porpoise | 974,967 | 739,024 | 0.138 | 130.60 |
| Pilotwhale | Whitesided | Finless | 861,276 | 855,083 | 0.004 | 4.41 |
| Pilotwhale | Whitesided | Harbour porpoise | 892,930 | 884,620 | 0.005 | 5.64 |
| Pilotwhale | Bottlenose | Finless | 828,193 | 724,397 | 0.067 | 73.75 |
| Pilotwhale | Bottlenose | Harbour porpoise | 857,823 | 749,827 | 0.067 | 76.38 |
| Pilotwhale | IndoBottlenose | Finless | 829,393 | 692,413 | 0.090 | 97.23 |
| Pilotwhale | IndoBottlenose | Harbour porpoise | 859,146 | 718,044 | 0.089 | 98.69 |
| Whitesided | Bottlenose | Harbour porpoise | 887,876 | 787,914 | 0.060 | 68.88 |
| Whitesided | Bottlenose | Finless | 857,483 | 760,224 | 0.060 | 69.75 |
| Whitesided | IndoBottlenose | Harbour porpoise | 888,872 | 755,955 | 0.081 | 92.25 |
| Whitesided | IndoBottlenose | Finless | 858,523 | 727,924 | 0.082 | 92.84 |
| Bottlenose | IndoBottlenose | Narwhal | 414,272 | 380,995 | 0.042 | 33.84 |
| Bottlenose | IndoBottlenose | Beluga | 434,366 | 396,566 | 0.045 | 37.67 |
| Killer whale | Pilotwhale | Narwhal | 955,756 | 837,598 | 0.066 | 61.58 |
| Killer whale | Pilotwhale | Beluga | 984,462 | 854,528 | 0.071 | 65.67 |
| Killer whale | Whitesided | Narwhal | 953,496 | 826,881 | 0.071 | 66.17 |
| Killer whale | Whitesided | Beluga | 982,162 | 844,661 | 0.075 | 67.95 |
| Killer whale | Bottlenose | Narwhal | 971,164 | 751,458 | 0.128 | 111.86 |
| Killer whale | Bottlenose | Beluga | 1,001,546 | 767,422 | 0.132 | 113.69 |
| Killer whale | IndoBottlenose | Narwhal | 974,507 | 722,249 | 0.149 | 126.51 |
| Killer whale | IndoBottlenose | Beluga | 1,007,582 | 736,424 | 0.155 | 128.87 |
| Pilotwhale | Whitesided | Beluga | 918,941 | 911,423 | 0.004 | 4.93 |
| Pilotwhale | Whitesided | Narwhal | 891,298 | 883,114 | 0.005 | 5.61 |
| Pilotwhale | Bottlenose | Narwhal | 859,652 | 743,735 | 0.072 | 78.60 |
| Pilotwhale | Bottlenose | Beluga | 887,196 | 766,562 | 0.073 | 81.55 |
| Pilotwhale | IndoBottlenose | Narwhal | 863,608 | 710,777 | 0.097 | 103.83 |
| Pilotwhale | IndoBottlenose | Beluga | 895,023 | 731,826 | 0.100 | 105.92 |
| Whitesided | Bottlenose | Narwhal | 888,390 | 780,573 | 0.065 | 74.77 |
| Whitesided | Bottlenose | Beluga | 917,400 | 804,237 | 0.066 | 76.44 |
| Whitesided | IndoBottlenose | Narwhal | 892,496 | 747,539 | 0.088 | 97.69 |
| Whitesided | IndoBottlenose | Beluga | 925,091 | 769,228 | 0.092 | 102.86 |

| | | | | | | |
|---|---|---|---|---|---|---|
| **Finless** | **Harbour porpoise** | **Narwhal** | **452,411** | **450,657** | **0.002** | **1.59** |
| Harbour porpoise | Finless | Beluga | 570,767 | 552,830 | 0.016 | 13.47 |
| Narwhal | Beluga | Harbour porpoise | 532,605 | 502,660 | 0.029 | 25.72 |
| Narwhal | Beluga | Finless | 514,273 | 466,273 | 0.049 | 41.75 |
| Finless | Narwhal | Killer whale | 973,140 | 885,678 | 0.047 | 47.30 |
| Finless | Narwhal | Bottlenose | 1,077,206 | 966,370 | 0.054 | 55.93 |
| Finless | Narwhal | IndoBottlenose | 1,080,812 | 970,600 | 0.054 | 56.63 |
| Finless | Narwhal | Pilotwhale | 1,059,846 | 950,178 | 0.055 | 57.27 |
| Finless | Beluga | Killer whale | 989,901 | 875,364 | 0.061 | 57.51 |
| Finless | Narwhal | Whitesided | 1,062,632 | 951,040 | 0.055 | 57.94 |
| Finless | Beluga | Bottlenose | 1,103,352 | 951,967 | 0.074 | 68.54 |
| Finless | Beluga | Pilotwhale | 1,084,679 | 936,511 | 0.073 | 68.84 |
| Finless | Beluga | IndoBottlenose | 1,109,158 | 955,589 | 0.074 | 69.72 |
| Finless | Beluga | Whitesided | 1,087,277 | 938,148 | 0.074 | 69.88 |
| Harbour porpoise | Narwhal | Killer whale | 1,004,793 | 891,909 | 0.060 | 59.43 |
| Harbour porpoise | Beluga | Killer whale | 1,028,676 | 885,849 | 0.075 | 69.85 |
| Harbour porpoise | Narwhal | Pilotwhale | 1,124,641 | 974,232 | 0.072 | 75.43 |
| Harbour porpoise | Narwhal | Bottlenose | 1,145,470 | 990,640 | 0.072 | 75.66 |
| Harbour porpoise | Narwhal | Whitesided | 1,127,578 | 976,951 | 0.072 | 75.84 |
| Harbour porpoise | Narwhal | IndoBottlenose | 1,153,263 | 994,022 | 0.074 | 78.93 |
| Harbour porpoise | Beluga | Pilotwhale | 1,163,136 | 965,266 | 0.093 | 88.73 |
| Harbour porpoise | Beluga | Whitesided | 1,165,862 | 968,086 | 0.093 | 89.42 |
| Harbour porpoise | Beluga | Bottlenose | 1,185,612 | 981,030 | 0.094 | 89.66 |
| Harbour porpoise | Beluga | IndoBottlenose | 1,197,547 | 984,311 | 0.098 | 93.10 |

**Supplementary table S6:** 100kb non-overlapping sliding window D-foil results for all quadruplet combinations [[H1,H2][H3,H4]] phylogenetically concurrent with our consensus topology shown in figure 1. Baiji was used as the outgroup/ancestral sequence. - attached as a spreadsheet. NA indicates not enough data in the window. None indicates no gene flow. As we implemented many different combinations, the species designation to H1 - H4 is indicated at the top of the table. Numbers within the table show the number of windows that show evidence to the gene flow event depicted. - attached as spreadsheet

**Supplementary table S7:** The pre-divergence Ne, divergence time intervals, and the increments specified for each of the species pair used for the simulations to compare against the hPSMC results.

| **Species pair** | **Pre-divergence Ne** | **Range (Ma)** | **Increments (years)** |
|---|---|---|---|
| Beluga whale + Narwhal | 30,000 | 1-2 | 100,000 |
| Beluga whale + Finless porpoise | 60,000 | 3-7 | 200,000 |
| Beluga whale + Harbour porpoise | 60,000 | 3-7 | 200,000 |
| Narwhal + Finless porpoise | 60,000 | 3-7 | 200,000 |
| Narwhal + Harbour porpoise | 60,000 | 3-7 | 200,000 |
| Beluga whale + Bottlenose dolphin | 105,000 | 3.9-8.5 | 200,000 |
| Beluga whale + Indo-Pacific bottlenose dolphin | 105,000 | 3.9-8.5 | 200,000 |
| Narwhal + Bottlenose dolphin | 105,000 | 3.9-8.5 | 200,000 |
| Narwhal + Indo-Pacific bottlenose dolphin | 105,000 | 3.9-8.5 | 200,000 |
| Narwhal + Killer whale | 105,000 | 3.9-8.5 | 200,000 |
| Narwhal + Long-finned pilot whale | 105,000 | 3.9-8.5 | 200,000 |
| Narwhal + Pacific white-sided dolphin | 105,000 | 3.9-8.5 | 200,000 |
| Beluga whale + Killer whale | 105,000 | 3.9-8.5 | 200,000 |
| Beluga whale + Long-finned pilot whale | 105,000 | 3.9-8.5 | 200,000 |
| Beluga whale + Pacific white-sided dolphin | 105,000 | 3.9-8.5 | 200,000 |
| Harbour porpoise + Bottlenose dolphin | 105,000 | 3.9-8.5 | 200,000 |
| Harbour porpoise + Indo-Pacific bottlenose dolphin | 105,000 | 3.9-8.5 | 200,000 |
| Finless porpoise + Bottlenose dolphin | 105,000 | 3.9-8.5 | 200,000 |
| Finless porpoise + Indo-Pacific bottlenose dolphin | 105,000 | 3.9-8.5 | 200,000 |
| Finless porpoise + Killer whale | 105,000 | 3.9-8.5 | 200,000 |
| Finless porpoise + Long-finned pilot whale | 105,000 | 3.9-8.5 | 200,000 |
| Finless porpoise + Pacific white-sided dolphin | 105,000 | 3.9-8.5 | 200,000 |
| Harbour porpoise + Killer whale | 105,000 | 3.9-8.5 | 200,000 |
| Harbour porpoise + Long-finned pilot whale | 105,000 | 3.9-8.5 | 200,000 |

| | | | |
|---|---|---|---|
| Harbour porpoise + Pacific white-sided dolphin | 105,000 | 3.9-8.5 | 200,000 |
| Harbour porpoise + Finless porpoise | 40,000 | 0.3-1.4 | 100,000 |
| Indo-Pacific Bottlenose dolphin dolphin + Bottlenose dolphin | 20,000 | 0.2-1.2 | 100,000 |
| Indo-Pacific bottlenose dolphin + Killer whale | 50,000 | 0.9-2.1 & 3.4-7 | 200,000 |
| Indo-Pacific bottlenose dolphin + Long-finned pilot whale | 50,000 | 0.9-2.1 & 3.4-7 | 200,000 |
| Indo-Pacific bottlenose dolphin + Pacific white-sided dolphin | 50,000 | 0.9-2.1 & 3.4-7 | 200,000 |
| Bottlenose dolphin + Killer whale | 50,000 | 0.9-2.1 & 3.4-7 | 200,000 |
| Bottlenose dolphin + Long-finned pilot whale | 50,000 | 0.9-2.1 & 3.4-7 | 200,000 |
| Bottlenose dolphin + Pacific white-sided dolphin | 50,000 | 0.9-2.1 & 3.4-7 | 200,000 |
| Long-finned pilot whale + Killer whale | 60,000 | 0.9-2.1 & 3.4-7 | 200,000 |
| Pacific white-sided dolphin + Killer whale | 50,000 | 0.9-2.1 & 3.4-7 | 200,000 |
| Pacific white-sided dolphin + Long-finned pilot whale | 50,000 | 0.9-2.1 & 3.4-7 | 200,000 |

**Supplementary table S8:** Mapping statistics of each Delphinoidea species used in this study when specifying the reference genome as the baiji assembly.

| Common name | Raw read pairs | Mapped reads | Coverage | Bp-mapped |
|---|---|---|---|---|
| Beluga | 466,374,135 | 476,814,543 | 31.44 | 69,807,010,359 |
| Bottlenose dolphin | 578,690,171 | 732,418,659 | 47.61 | 105,524,983,813 |
| Harbour porpoise | 289,063,910 | 418,431,029 | 23.17 | 50,830,083,145 |
| Indo-Pacific bottlenose dolphin | 466,306,082 | 551,837,703 | 35.62 | 78,749,625,267 |
| Indo-Pacific finless porpoise | 523,612,238 | 557,766,873 | 24.96 | 54,450,935,944 |
| Killer whale | 1,467,089,287 | 1,047,260,000 | 39.53 | 88,692,400,000 |
| Long-finned pilot whale | 428,064,233 | 504,482,080 | 28.61 | 63,276,638,573 |
| Narwhal | 384,563,392 | 468,429,237 | 31.09 | 68,247,058,370 |
| Pacific white-sided dolphin | 453,348,710 | 499,704,592 | 28.83 | 63,800,396,300 |

**Supplementary table S9:** Mapping statistics of each Delphinoidea species used in this study when specifying the reference genome as a conspecific assembly.

| Common name | Raw read pairs | Mapped reads | Coverage | Bp-mapped |
|---|---|---|---|---|
| Beluga | 466,374,135 | 531,535,936 | 34.47 | 79,218,898,913 |
| Bottlenose dolphin | 578,690,171 | 779,210,277 | 54.03 | 114,530,169,747 |
| Harbour porpoise | 289,063,910 | 431,762,883 | 23.74 | 52,067,455,809 |
| Indo-Pacific bottlenose dolphin | 466,306,082 | 587,440,922 | 37.88 | 85,032,333,848 |
| Indo-Pacific finless porpoise | 523,612,238 | 620,580,505 | 27.33 | 61,286,732,910 |
| Killer whale | 1,467,089,287 | 1,213,221,913 | 44.93 | 100,903,316,971 |
| Long-finned pilot whale | 428,064,233 | 598,612,204 | 32.79 | 75,639,560,432 |
| Narwhal | 384,563,392 | 529,082,769 | 33.85 | 78,238,763,386 |
| Pacific white-sided dolphin | 453,348,710 | 592,814,373 | 33.02 | 76,299,243,217 |

**Supplementary table S10:** Genome-wide pairwise distance matrix of the nine Delphinoidea included in this study. Bottlenose = bottlenose dolphin, Finless = finless porpoise, Harbour = harbour porpoise, Indobottle = Indo-Pacfic bottlenose dolphin, Killer = killer whale, Pilot = pilot whale, White = Pacific white-sided dolphin.

| | | | | | | | | | |
|---|---|---|---|---|---|---|---|---|---|
| Beluga | 0.0000 | 0.0211 | 0.0151 | 0.0153 | 0.0211 | 0.0205 | 0.0056 | 0.0210 | 0.0209 |
| Bottlenose | 0.0211 | 0.0000 | 0.0230 | 0.0231 | 0.0040 | 0.0113 | 0.0210 | 0.0102 | 0.0107 |
| Finless | 0.0151 | 0.0230 | 0.0000 | 0.0056 | 0.0230 | 0.0224 | 0.0151 | 0.0229 | 0.0228 |
| Harbour | 0.0153 | 0.0231 | 0.0056 | 0.0000 | 0.0231 | 0.0225 | 0.0152 | 0.0231 | 0.0230 |
| Indobottle | 0.0211 | 0.0040 | 0.0230 | 0.0231 | 0.0000 | 0.0113 | 0.0210 | 0.0102 | 0.0107 |
| Killer | 0.0205 | 0.0113 | 0.0224 | 0.0225 | 0.0113 | 0.0000 | 0.0204 | 0.0113 | 0.0112 |
| Narwhal | 0.0056 | 0.0210 | 0.0151 | 0.0152 | 0.0210 | 0.0204 | 0.0000 | 0.0209 | 0.0208 |
| Pilot | 0.0210 | 0.0102 | 0.0229 | 0.0231 | 0.0102 | 0.0113 | 0.0209 | 0.0000 | 0.0109 |
| White | 0.0209 | 0.0107 | 0.0228 | 0.0230 | 0.0107 | 0.0112 | 0.0208 | 0.0109 | 0.0000 |

**Supplementary table S11:** Metrics used to calculate the mutation rate per year with the equation mutation rate = divergence time / 2x genetic distance. Mean divergences were taken from the full dataset 10-partition AR from McGowen et al 2020 (McGowen et al., 2020) and average genetic distances were calculated from the results shown in supplementary table S5.

| **Species** | **Closest relative** | **Divergence (Ma)** | **Distance** | **Mutation rate per year** |
|---|---|---|---|---|
| Beluga | Narwhal | 7.72 | 0.0056 | $3.63\text{x}10^{-10}$ |
| Killer whale | Delphinidae | 10.16 | 0.0113 | $5.56\text{x}10^{-10}$ |
| Bottlenose dolphin | Indo-Pacific bottlenose dolphin | 2.69 | 0.0040 | $7.51\text{x}10^{-10}$ |
| Harbour porpoise | Finless porpoise | 5.36 | 0.0056 | $5.25\text{x}10^{-10}$ |
| Long-finned pilot whale | *Tursiops* spp. | 7.46 | 0.0102 | $6.83\text{x}10^{-10}$ |
| Pacific while-sided dolphin | *Tursiops* + *Globicephala* | 9.48 | 0.0108 | $5.69\text{x}10^{-10}$ |

**Supplementary table S12**: Generation times, generational mutation rates and references for the generation times for each of the nine Delphinoidea species used in this study.

| **Common name** | **Generation time** | **Generational mutation rate** | **Generation time reference** | **Bp-mapped** |
|---|---|---|---|---|
| Beluga | 32 | $1.16x10^{-8}$ | (Garde et al., 2015) | 79,218,898,913 |
| Bottlenose dolphin | 21 | $1.58x10^{-8}$ | (Taylor et al., 2007) | 114,530,169,747 |
| Harbour porpoise | 10 | $5.25x10^{-9}$ | (Birkun and Frantzis, 2008) | 52,067,455,809 |
| Indo-Pacific bottlenose dolphin | 21 | $1.58x10^{-8}$ | (Taylor et al., 2007) | 85,032,333,848 |
| Indo-Pacific finless porpoise | 8 | $4.20x10^{-9}$ | (Zhou et al., 2018) | 61,286,732,910 |
| Killer whale | 26 | $1.43x10^{-8}$ | (Foote et al., 2016) | 100,903,316,971 |
| Long-finned pilot whale | 24 | $1.64x10^{-8}$ | (Taylor et al., 2007) | 75,639,560,432 |
| Narwhal | 30 | $1.09x10^{-8}$ | (Garde et al., 2015) | 78,238,763,386 |
| Pacific white-sided dolphin | 21 | $1.21x10^{-8}$ | (Taylor et al., 2007) | 76,299,243,217 |

### Supplementary figures

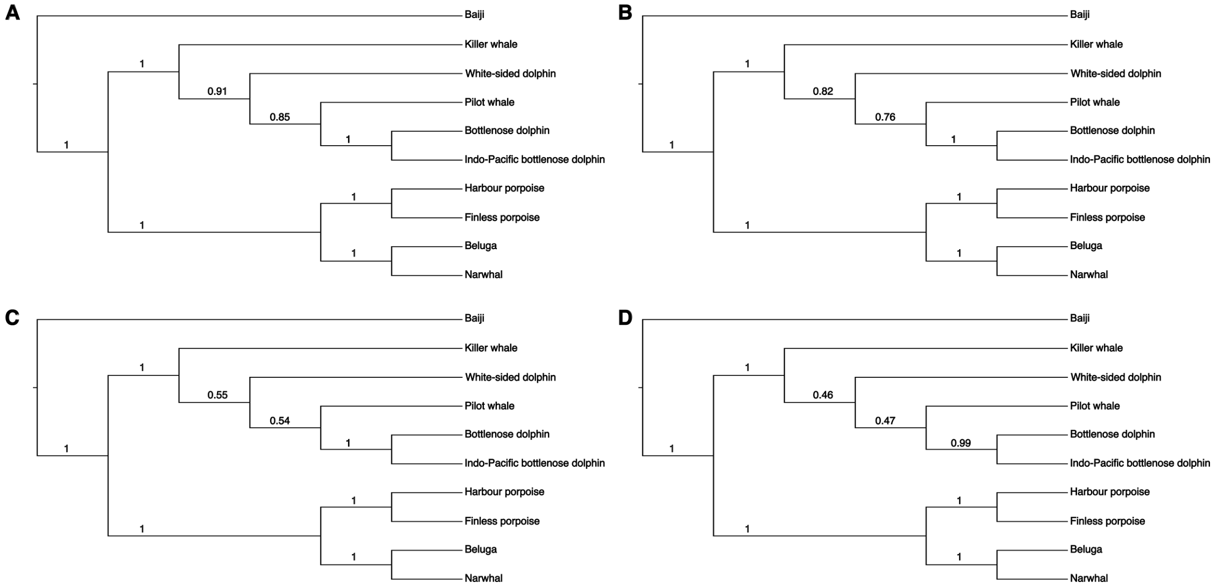


**Supplementary figure S1:** Consensus trees of independent Maximum-Likelihood trees constructed from non-overlapping sliding windows of (A) 1Mb, (B) 500kb, (C) 100kb, or (D) 50kb in length. Branch numbers represent the number of independent trees supporting each node.

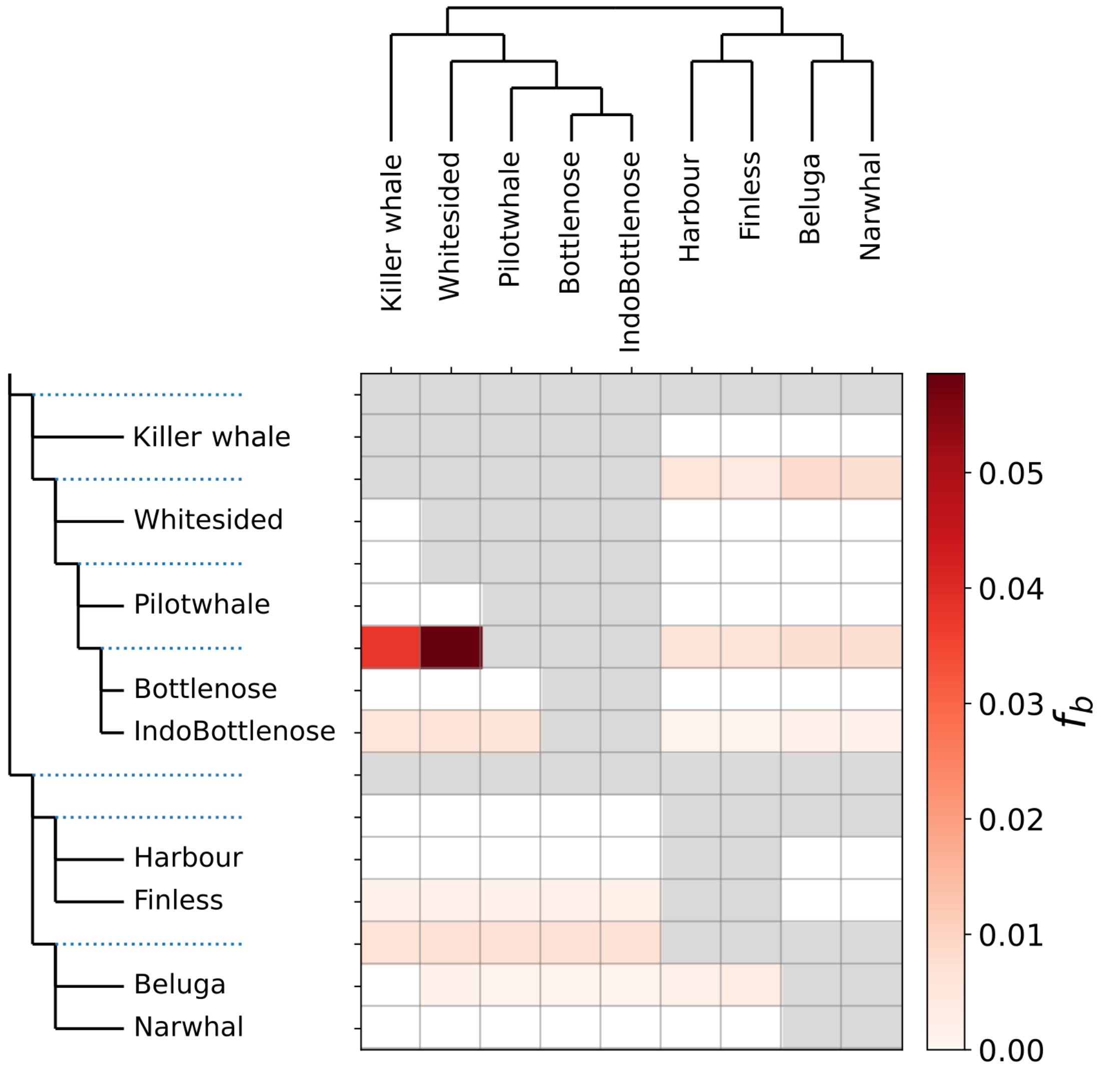


**Supplementary figure S2:** X chromosome Fbranch results. The species tree is displayed above while the trees to the left and write of the matrix are an expanded form, including internal branches as dotted lines. The values in the matrix refer to excess allele sharing between the expanded tree branch (relative to its sister branch) and the species on the *x*-axis.

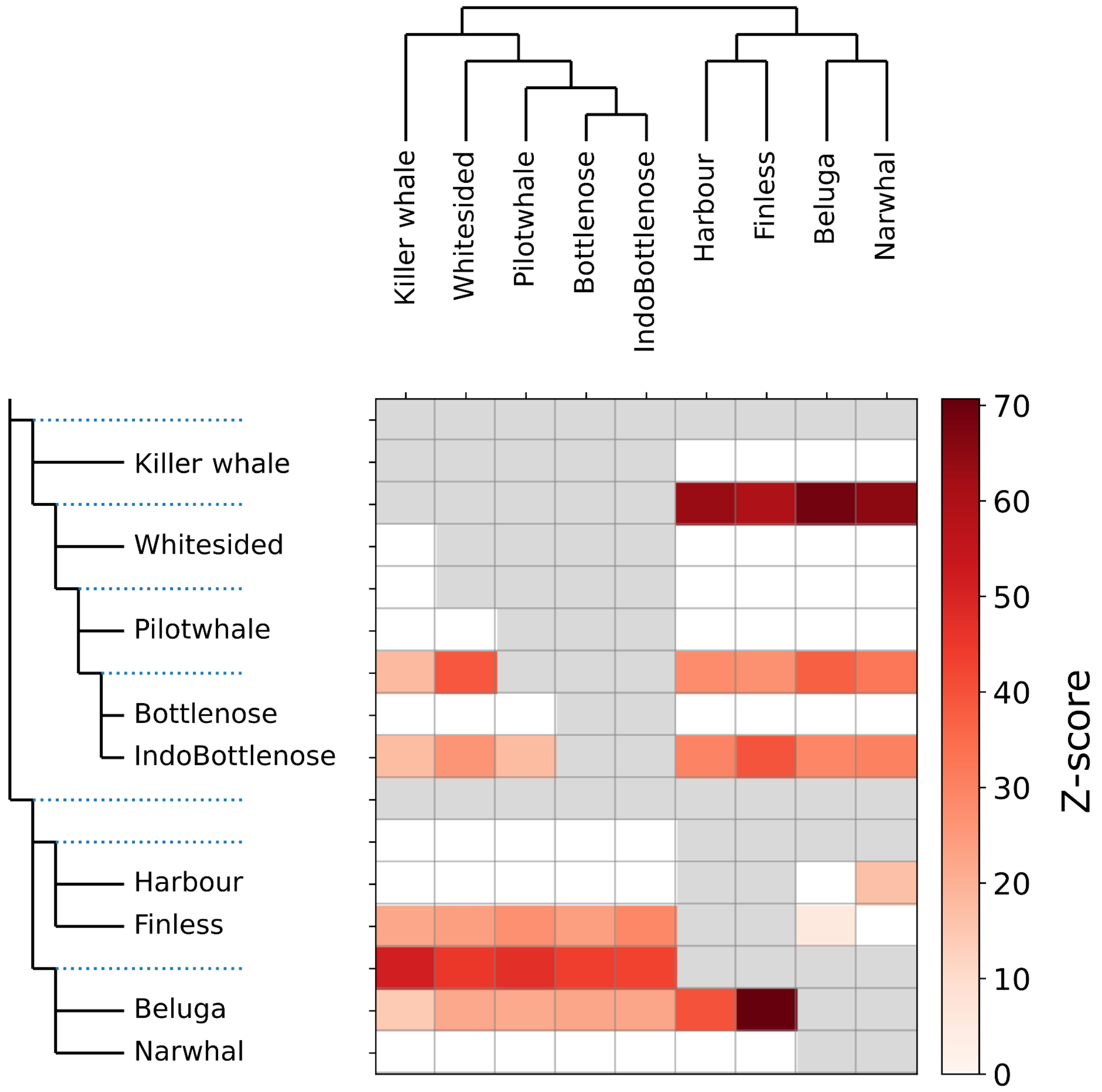


**Supplementary figure S3:** Significance (Z-score) of the autosomal *f*-branch results. A |Z|>3 is considered significant. The values in the matrix refer to Z-score for the *f*b value (Fig 2) between the expanded tree branch (relative to its sister branch) and the species on the *x*-axis.

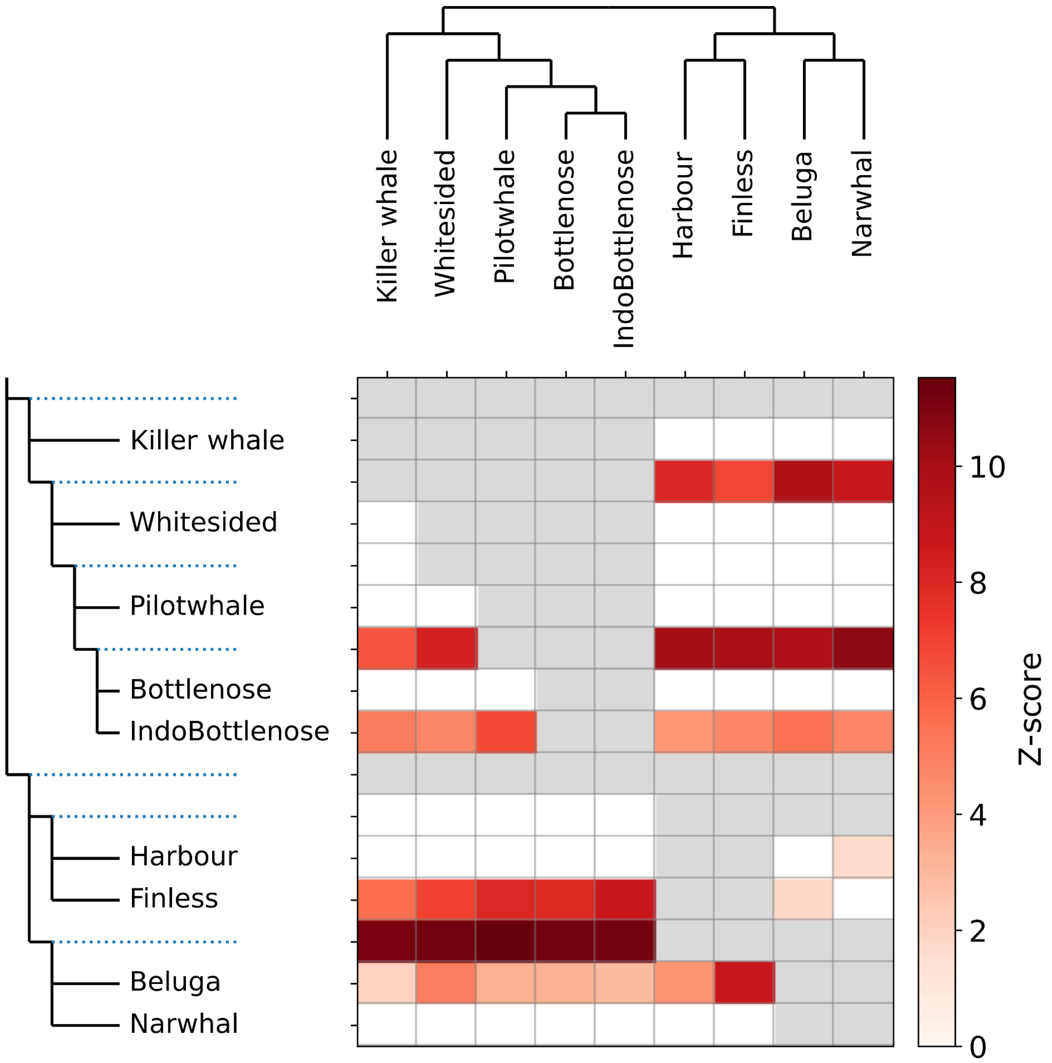


**Supplementary figure S4:** Significance (Z-score) of the X chromosome *f*-branch results. A |Z|>3 is considered significant. The values in the matrix refer to Z-score for the *f*b value (Supplementary Fig S2) between the expanded tree branch (relative to its sister branch) and the species on the *x*-axis.

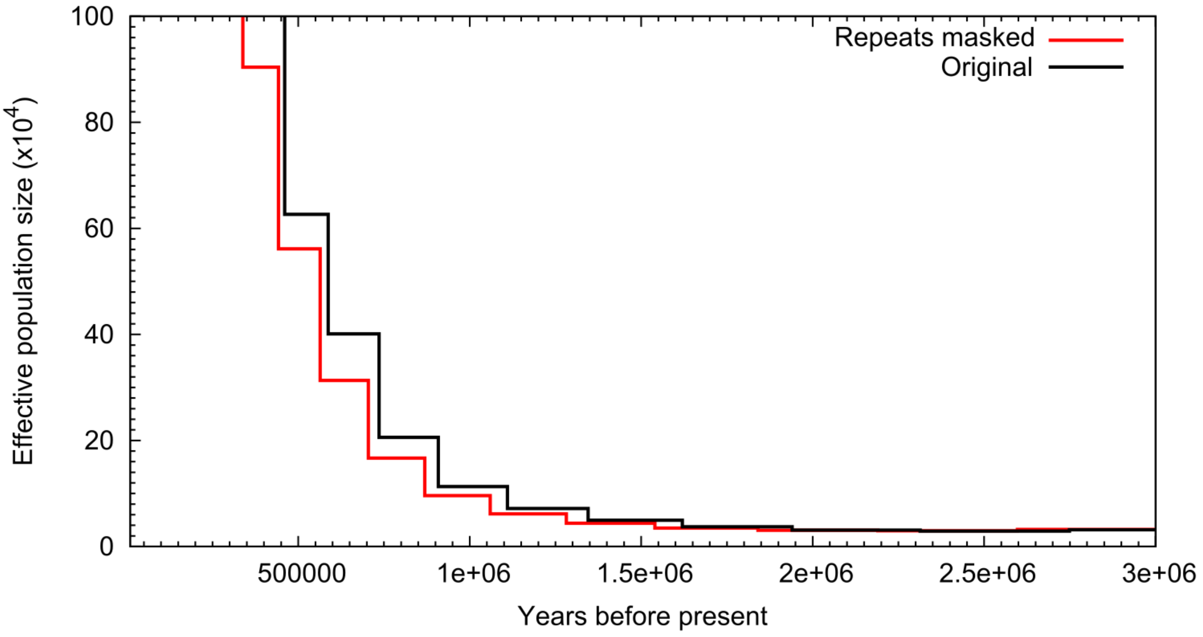


**Supplementary figure S5:** Comparison of hPSMC results using a pseudodiploid sequence from the bottlenose and Indo-Pacific bottlenose dolphins (shallow divergence) with either repeat regions masked or not.

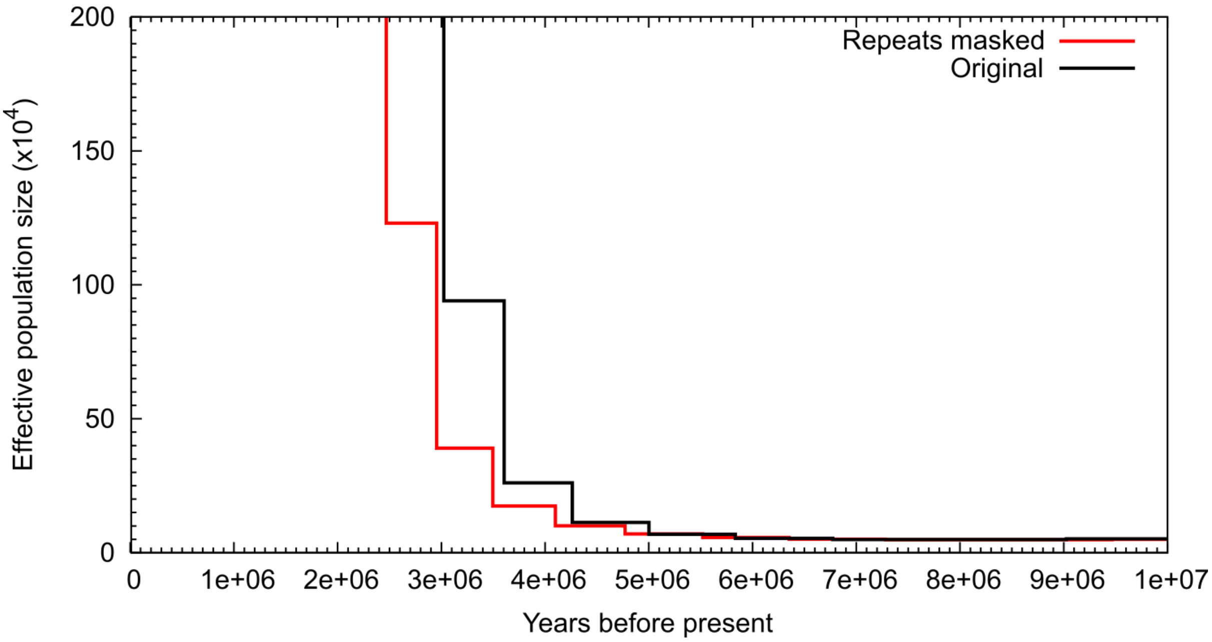


**Supplementary figure S6:** Comparison of hPSMC results using a pseudodiploid sequence from the beluga and narwhal (medium divergence) with either repeat regions masked or not.

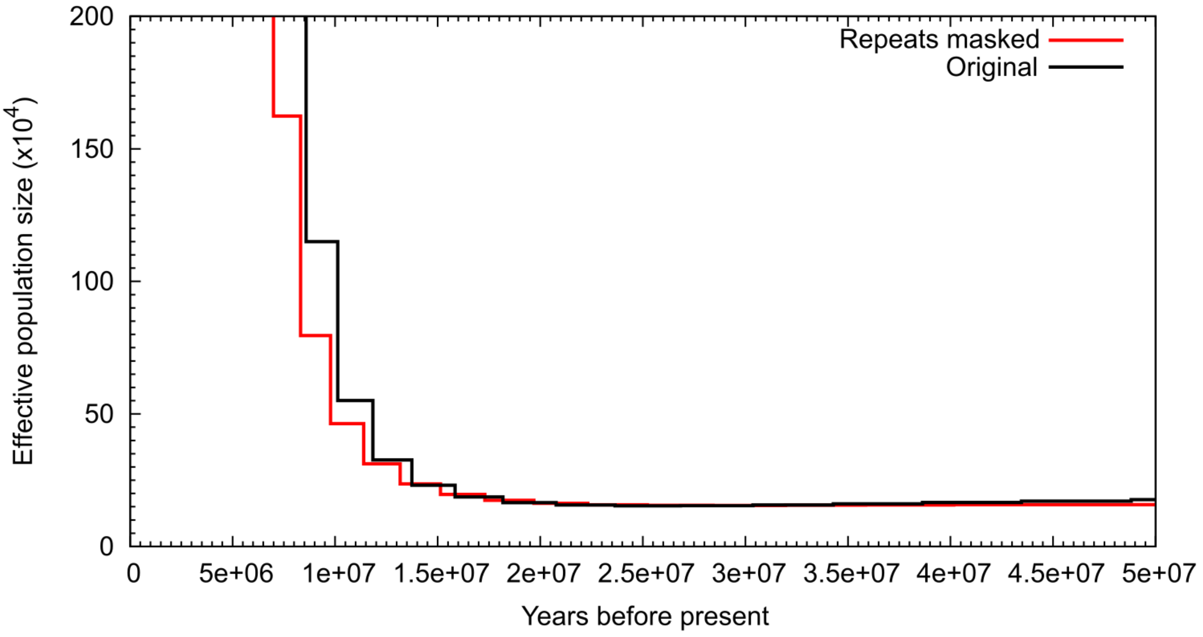


**Supplementary figure S7:** Comparison of hPSMC results using a pseudodiploid sequence from the bottlenose dolphin and beluga (deep divergence) with either repeat regions masked or not.

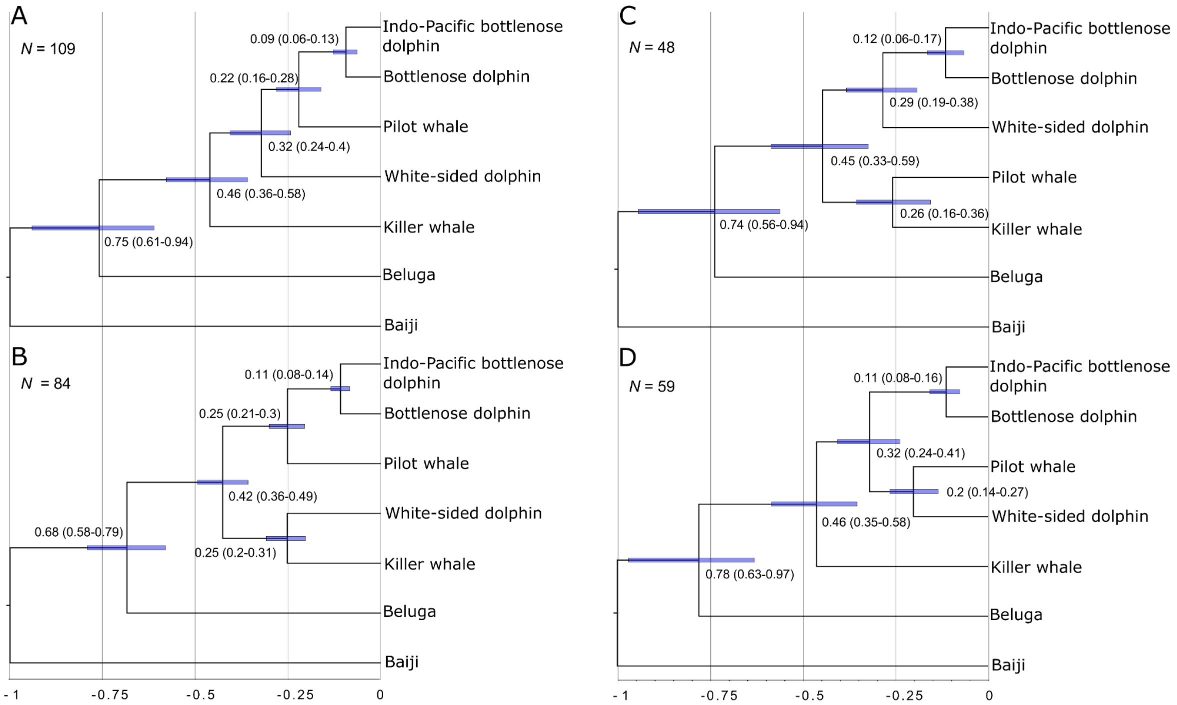


**Supplementary figure S8:** Relative divergence times of alternative topologies assumed to arise due to incomplete lineage sorting (ILS) or gene flow. N represents the number of independent loci supporting said topology. A) Consensus species topology. B) ILS/gene flow between the killer whale and Pacific white-sided dolphin. C) ILS/gene flow between killer whale and long-finned pilot whale. D) ILS/gene flow between Pacific white-sided dolphin and the long-finned pilot whale. Blue bars and numbers in parentheses show 95% credibility intervals.

### Supplementary results - hPSMC

Additional plots of the hPSMC empirical and simulated data can be found under the following link: https://sid.erda.dk/cgi-sid/ls.py?share_id=ewvczfS2hH on the University of Copenhagen's electronic research data archive (ERDA). Bold lines show the hPSMC empirical data, faded lines show the simulated data, and the black lines show the simulated data that most closely match the empirical data without overlapping it between 1.5x and 10x the pre-divergence Ne.

**Supplementary references**

Birkun AA Jr, Frantzis A. 2008. *Phocoena phocoena* ssp. relicta. The IUCN Red List of Threatened Species : e.T17027A50369903. https://dx.doi.org/10.2305/IUCN.UK.2020-2.RLTS.T17027A50369903.en.

Foote AD, Vijay N, Ávila-Arcos MC, Baird RW, Durban JW, Fumagalli M, Gibbs RA, Hanson MB, Korneliussen TS, Martin MD, Robertson KM, Sousa VC, Vieira FG, Vinař T, Wade P, Worley KC, Excoffier L, Morin PA, Gilbert MTP, Wolf JBW. 2016. Genome-culture coevolution promotes rapid divergence of killer whale ecotypes. *Nat Commun* **7**:11693.

Garde E, Hansen SH, Ditlevsen S, Tvermosegaard KB, Hansen J, Harding KC, Heide-Jørgensen MP. 2015. Life history parameters of narwhals (*Monodon monoceros*) from Greenland. *J Mammal* **96**:866–879.

McGowen MR, Tsagkogeorga G, Álvarez-Carretero S, Dos Reis M, Struebig M, Deaville R, Jepson PD, Jarman S, Polanowski A, Morin PA, Rossiter SJ. 2020. Phylogenomic Resolution of the Cetacean Tree of Life Using Target Sequence Capture. *Syst Biol* **69**:479–501.

Taylor BL, Chivers SJ, Larese J, Perrin WF. 2007. Generation length and percent mature estimates for IUCN assessments of cetaceans (No. Administrative Report LJ-07-01 ). National Marine Fisheries Service, Southwest Fisheries Science Center.

Zhou X, Guang X, Sun D, Xu S, Li M, Seim I, Jie W, Yang L, Zhu Q, Xu J, Gao Q, Kaya A, Dou Q, Chen B, Ren W, Li S, Zhou K, Gladyshev VN, Nielsen R, Fang X, Yang G. 2018. Population genomics of finless porpoises reveal an incipient cetacean species adapted to freshwater. *Nat Commun* **9**:1276.
