## Supplementary information for "A genomic assessment of the marine-speciation paradox within the toothed whale superfamily Delphinoidea" (from another version of this preprint: version 6, 2022-03-28)

| Triplet analysed | Geneflow pair | Control taxon | BIC2Dist (IBS + Geneflow) | BIC1Dist (IBS alone) | BIC difference | Significant for gene flow (BIC difference >10) | Number of trees | Percentage of total trees (2161) from triplet | Percentage of trees supporting topology expained by gene flow |
|---|---|---|---|---|---|---|---|---|---|
| Pilot whale_Bottlenose dolphin_Killer whale | Bot-Orca | Pilot whale | -4176.75 | -4015.52 | -161.23 | Yes | 363 | 16.80 | 44.13 |
| White-sided dolphin_Bottlenose dolphin_Killer whale | Bot-Orca | White-sided dolphin | -5203 | -5001.75 | -201.25 | Yes | 451 | 20.87 | 51.55 |
| Pilot whale_Indo-Pacific Bottlenose dolphin_Killer wha | Indo-Orca | Pilot whale | -4163.39 | -4003.35 | -160.04 | Yes | 362 | 16.75 | 44.27 |
| White-sided dolphin_Indo-Pacific Bottlenose dolphin_ | Indo-Orca | White-sided dolphin | -5157.77 | -4961.79 | -195.98 | Yes | 448 | 20.73 | 91.82 |
| Pilot whale_Bottlenose dolphin_Killer whale | Pilot-Orca | Bottlenose dolphin | -4149.09 | -3995.26 | -153.83 | Yes | 353 | 16.34 | 26.63 |
| Pilot whale_Indo-Pacific Bottlenose dolphin_Killer wha | Pilot-Orca | Indo-Pacific Bottlenose | -4145.01 | -3991.4 | -153.61 | Yes | 353 | 16.34 | 24.46 |
| White-sided dolphin_Pilot whale_Killer whale | Pilot-Orca | White-sided dolphin | -5551.99 | -5354.47 | -197.52 | Yes | 479 | 22.17 | 30.52 |
| White-sided dolphin_Pilot whale_Bottlenose dolphin | Pilot-White | Bottlenose dolphin | -5329.17 | -5126.07 | -203.10 | Yes | 459 | 21.24 | 44.05 |
| White-sided dolphin_Pilot whale_Indo-Pacific Bottlenc | Pilot-White | Indo-Pacific Bottlenose | -5332.08 | -5127.41 | -204.67 | Yes | 459 | 21.24 | 37.09 |
| White-sided dolphin_Pilot whale_Bottlenose dolphin | White-Bot | Pilot whale | -7160.67 | -6929.73 | -230.94 | Yes | 629 | 29.11 | 86.33 |
| White-sided dolphin_Pilot whale_Indo-Pacific Bottlenc | White-Indo | Pilot whale | -7154.12 | -6919.18 | -234.94 | Yes | 628 | 29.06 | 49.33 |
| White-sided dolphin_Bottlenose dolphin_Killer whale | White-Orca | Bottlenose dolphin | -5679.95 | -5365.25 | -314.70 | Yes | 478 | 22.12 | 29.40 |
| White-sided dolphin_Indo-Pacific Bottlenose dolphin_ | White-Orca | Indo-Pacific Bottlenose | -5687.27 | -5373.09 | -314.18 | Yes | 479 | 22.17 | 31.43 |
| White-sided dolphin_Pilot whale_Killer whale | White-Orca | Pilot whale | -6205.88 | -5910.93 | -294.95 | Yes | 529 | 24.48 | 50.04 |
| Indo-Pacific Bottlenose dolphin_Bottlenose dolphin_Ki | Bot-Orca | Indo-Pacific Bottlenose | -47.1718 | -40.833 | -6.34 | No | 4 | 0.19 | 1.09 |
| Indo-Pacific Bottlenose dolphin_Bottlenose dolphin_Ki | Indo-Orca | Bottlenose dolphin | -35.0559 | -32.055 | -3.00 | No | 3 | 0.14 | 0.37 |
| Pilot whale_Indo-Pacific Bottlenose dolphin_Bottlenos | Pilot-Bot | Indo-Pacific Bottlenose | -56.1656 | -53.3674 | -2.80 | No | 5 | 0.23 | 1.09 |
| Pilot whale_Indo-Pacific Bottlenose dolphin_Bottlenos | Pilot-Indo | Bottlenose dolphin | -43.6088 | -44.5198 | 0.91 | No | 4 | 0.19 | 0.15 |
| White-sided dolphin_Indo-Pacific Bottlenose dolphin_ | White-Bot | Indo-Pacific Bottlenose | -53.2849 | -53.8868 | 0.60 | No | 5 | 0.23 | 0.46 |
| White-sided dolphin_Indo-Pacific Bottlenose dolphin_ | White-Indo | Bottlenose dolphin | -41.6525 | -42.3186 | 0.67 | No | 4 | 0.19 | 0.31 |

| Triplet analysed | Geneflow pair | Control taxon | BIC2Dist (IBS + Geneflow) | BIC1Dist (IBS alone) | BIC difference | Significant for gene flow | Number of trees | % of total trees | % of trees supporting topology expained by gene flow |
|---|---|---|---|---|---|---|---|---|---|
| Pilot whale_Bottlenose dolphin_Killer whale | Bot-Orca | Pilot whale | -5877.09 | -5828.01 | -49.08 | Yes | 543 | 19.89 | 12.79 |
| White-sided dolphin_Bottlenose dolphin_Killer whale | Bot-Orca | White-sided dolphin | -6493.50 | -6410.93 | -82.57 | Yes | 589 | 21.58 | 14.76 |
| Pilot whale_Indo-Pacific Bottlenose dolphin_Killer whale | Indo-Orca | Pilot whale | -5836.61 | -5777.56 | -59.05 | Yes | 539 | 19.74 | 13.24 |
| White-sided dolphin_Indo-Pacific Bottlenose dolphin_Killer whale | Indo-Orca | White-sided dolphin | -6501.26 | -6417.36 | -83.90 | Yes | 590 | 21.61 | 14.82 |
| Pilot whale_White-sided dolphin_Killer whale | Pilot-Orca | White-sided dolphin | -6892.35 | -6861.90 | -30.45 | Yes | 631 | 23.11 | 12.75 |
| Pilot whale_White-sided dolphin_Bottlenose dolphin | Pilot-White | Bottlenose dolphin | -7033.39 | -6989.18 | -44.21 | Yes | 648 | 23.74 | 14.00 |
| Pilot whale_White-sided dolphin_Indo-Pacific Bottlenose dolphin | Pilot-White | Indo-Pacific Bottlenose dolphir | -7073.33 | -7026.60 | -46.73 | Yes | 651 | 23.85 | 14.15 |
| Pilot whale_White-sided dolphin_Bottlenose dolphin | White-Bot | Pilot whale | -9197.44 | -9186.93 | -10.51 | Yes | 865 | 31.68 | 16.05 |
| Pilot whale_White-sided dolphin_Killer whale | White-Orca | Pilot whale | -8498.20 | -8408.06 | -90.14 | Yes | 784 | 28.72 | 19.25 |
| White-sided dolphin_Bottlenose dolphin_Killer whale | White-Orca | Bottlenose dolphin | -7986.93 | -7853.23 | -133.70 | Yes | 726 | 26.59 | 19.83 |
| White-sided dolphin_Indo-Pacific Bottlenose dolphin_Killer whale | White-Orca | Indo-Pacific Bottlenose dolphir | -7983.67 | -7846.07 | -137.60 | Yes | 726 | 26.59 | 20.03 |
| Indo-Pacific Bottlenose dolphin_Bottlenose dolphin_Killer whale | Bot-Orca | Indo-Pacific Bottlenose dolphir | -143.55 | -144.83 | 1.28 | No | 13 | 0.48 | 0.39 |
| Indo-Pacific Bottlenose dolphin_Bottlenose dolphin_Killer whale | Indo-Orca | Bottlenose dolphin | -82.72 | -81.61 | -1.11 | No | 8 | 0.29 | 0.25 |
| Pilot whale_Indo-Pacific Bottlenose dolphin_Bottlenose dolphin | Pilot-Bot | Indo-Pacific Bottlenose dolphir | -306.79 | -305.15 | -1.64 | No | 28 | 1.03 | 0.82 |
| Pilot whale_Indo-Pacific Bottlenose dolphin_Bottlenose dolphin | Pilot-Indo | Bottlenose dolphin | -330.52 | -336.87 | 6.35 | No | 31 | 1.14 | 0.52 |
| Pilot whale_Bottlenose dolphin_Killer whale | Pilot-Orca | Bottlenose dolphin | -5643.28 | -5648.29 | 5.01 | No | 521 | 19.08 | 9.13 |
| Pilot whale_Indo-Pacific Bottlenose dolphin_Killer whale | Pilot-Orca | Indo-Pacific Bottlenose dolphir | -5701.86 | -5699.31 | -2.55 | No | 525 | 19.23 | 9.77 |
| White-sided dolphin_Indo-Pacific Bottlenose dolphin_Bottlenose d | White-Bot | Indo-Pacific Bottlenose dolphir | -257.04 | -258.27 | 1.24 | No | 24 | 0.88 | 0.56 |
| Pilot whale_White-sided dolphin_Indo-Pacific Bottlenose dolphin | White-Indo | Pilot whale | -9117.94 | -9115.68 | -2.26 | No | 858 | 31.43 | 15.41 |
| White-sided dolphin_Indo-Pacific Bottlenose dolphin_Bottlenose d | White-Indo | Bottlenose dolphin | -170.67 | -176.49 | 5.81 | No | 16 | 0.59 | 0.23 |

**Monodontidae vs Delphinidae**

| | | | | | | | | | | |
|---|---|---|---|---|---|---|---|---|---|---|
| **H1** | Beluga | Beluga | Beluga | Beluga | Beluga | Beluga | Beluga | Beluga | Beluga | Bottlenose dolphin |
| **H2** | Narwhal | Narwhal | Narwhal | Narwhal | Narwhal | Narwhal | Narwhal | Narwhal | Narwhal | Indo-Pacific Bottlenose dolphin |
| **H3** | Bottlenose dolphin | Indo-Pacific Bottlenose dolphin | Bottlenose dolphin | Indo-Pacific Bottlenose dolphin | Bottlenose dolphin | Indo-Pacific Bottlenose dolphin | Pilot whale | Pilot whale | White-sided dolphin | Beluga |
| **H4** | Killer whale | Killer whale | Pilot whale | Pilot whale | White-sided dolphin | White-sided dolphin | White-sided dolphin | Killer whale | Killer whale | Narwhal |
| **Gene flow from H1 into H3** | 1 | 0 | 0 | 2 | 0 | 0 | 0 | 1 | 1 | 0 |
| **Gene flow from H1 into H4** | 3 | 4 | 0 | 0 | 2 | 2 | 1 | 6 | 5 | 1 |
| **Gene flow from H2 into H3** | 0 | 0 | 0 | 2 | 0 | 0 | 1 | 0 | 0 | 0 |
| **Gene flow from H2 into H4** | 17 | 22 | 3 | 1 | 1 | 1 | 4 | 22 | 12 | 1 |
| **Gene flow from H3 into H1** | 1 | 0 | 0 | 0 | 0 | 1 | 0 | 0 | 0 | 0 |
| **Gene flow from H3 into H2** | 0 | 0 | 0 | 0 | 0 | 0 | 0 | 0 | 0 | 0 |
| **Gene flow from H4 into H1** | 3 | 2 | 0 | 0 | 0 | 1 | 0 | 3 | 0 | 0 |
| **Gene flow from H4 into H2** | 7 | 5 | 3 | 0 | 3 | 2 | 1 | 5 | 6 | 1 |
| **Gene flow between ancestor H1-H2 and H3** | 9 | 11 | 98 | 118 | 64 | 68 | 115 | 10 | 13 | 129 |
| **Gene flow between ancestor H1-H2 and H4** | 3480 | 3416 | 243 | 264 | 470 | 459 | 315 | 2686 | 1950 | 452 |
| **NA** | 7 | 7 | 10 | 11 | 8 | 10 | 10 | 10 | 8 | 10 |
| **None** | 18019 | 18080 | 21190 | 21149 | 20999 | 21003 | 21099 | 18804 | 19552 | 20953 |

**Phocoenidae vs Delphinidae**

| | | | | | | | | | | |
|---|---|---|---|---|---|---|---|---|---|---|
| **H1** | Finless | Finless | Finless | Finless | Finless | Finless | Finless | Finless | Finless | Bottlenose dolphin |
| **H2** | Harbour | Harbour | Harbour | Harbour | Harbour | Harbour | Harbour | Harbour | Harbour | Indo-Pacific Bottlenose dolphin |
| **H3** | Bottlenose dolphin | Indo-Pacific Bottlenose dolphin | Bottlenose dolphin | Indo-Pacific Bottlenose dolphin | Bottlenose dolphin | Indo-Pacific Bottlenose dolphin | Pilot whale | Pilot whale | White-sided dolphin | Finless |
| **H4** | Killer whale | Killer whale | Pilot whale | Pilot whale | White-sided dolphin | White-sided dolphin | White-sided dolphin | Killer whale | Killer whale | Harbour |
| **Gene flow from H1 into H3** | 0 | 0 | 0 | 1 | 0 | 0 | 0 | 0 | 0 | 0 |
| **Gene flow from H1 into H4** | 31 | 31 | 1 | 1 | 4 | 6 | 1 | 19 | 18 | 1 |
| **Gene flow from H2 into H3** | 0 | 0 | 0 | 0 | 0 | 0 | 0 | 0 | 0 | 1 |
| **Gene flow from H2 into H4** | 3 | 4 | 0 | 0 | 0 | 0 | 0 | 0 | 1 | 0 |
| **Gene flow from H3 into H1** | 1 | 0 | 1 | 0 | 1 | 1 | 0 | 0 | 1 | 0 |
| **Gene flow from H3 into H2** | 0 | 0 | 0 | 0 | 0 | 0 | 0 | 0 | 0 | 1 |
| **Gene flow from H4 into H1** | 11 | 8 | 1 | 3 | 5 | 1 | 1 | 10 | 9 | 0 |
| **Gene flow from H4 into H2** | 3 | 1 | 0 | 0 | 0 | 0 | 0 | 0 | 0 | 0 |
| **Gene flow between ancestor H1-H2 and H3** | 9 | 10 | 114 | 127 | 78 | 62 | 110 | 11 | 17 | 588 |
| **Gene flow between ancestor H1-H2 and H4** | 3284 | 3182 | 241 | 264 | 450 | 429 | 301 | 2521 | 1871 | 46 |
| **NA** | 11 | 13 | 12 | 12 | 14 | 14 | 15 | 13 | 11 | 18 |
| **None** | 18188 | 18292 | 21170 | 21132 | 20989 | 21029 | 21114 | 18967 | 19613 | 20886 |

**Phocoenidae vs Monodontidae**

| | | |
|---|---|---|
| **H1** | Beluga | Harbour |
| **H2** | Narwhal | Finless |
| **H3** | Harbour | Beluga |
| **H4** | Finless | Narwhal |
| **Gene flow from H1 into H3** | 1 | 2 |
| **Gene flow from H1 into H4** | 0 | 4 |
| **Gene flow from H2 into H3** | 0 | 0 |
| **Gene flow from H2 into H4** | 4 | 0 |
| **Gene flow from H3 into H1** | 0 | 0 |
| **Gene flow from H3 into H2** | 0 | 1 |
| **Gene flow from H4 into H1** | 2 | 4 |
| **Gene flow from H4 into H2** | 4 | 0 |
| **Gene flow between ancestor H1-H2 and H3** | 44 | 99 |
| **Gene flow between ancestor H1-H2 and H4** | 578 | 433 |
| **NA** | 10 | 10 |
| **None** | 20899 | 20989 |

**Delphinidae vs Monodontidae+Phocoenidae**

| | | | | | | | | | | |
|---|---|---|---|---|---|---|---|---|---|---|
| **H1** | White-sided dolphin | White-sided dolphin | White-sided dolphin | White-sided dolphin | Pilot whale | Pilot whale | Pilot whale | Pilot whale | Bottlenose dolphin | Bottlenose dolphin |
| **H2** | Killer whale | Killer whale | Killer whale | Killer whale | Bottlenose dolphin | Bottlenose dolphin | Bottlenose dolphin | Bottlenose dolphin | White-sided dolphin | White-sided dolphin |
| **H3** | Narwhal | Narwhal | Beluga | Beluga | Narwhal | Narwhal | Beluga | Beluga | Narwhal | Narwhal |
| **H4** | Harbour | Finless | Harbour | Finless | Harbour | Finless | Harbour | Finless | Harbour | Finless |
| **Gene flow from H1 into H3** | 1 | 1 | 4 | 1 | 69 | 73 | 72 | 66 | 30 | 25 |
| **Gene flow from H1 into H4** | 0 | 0 | 0 | 0 | 0 | 0 | 0 | 0 | 0 | 0 |
| **Gene flow from H2 into H3** | 368 | 313 | 362 | 335 | 24 | 24 | 28 | 28 | 119 | 119 |
| **Gene flow from H2 into H4** | 0 | 0 | 0 | 0 | 0 | 0 | 0 | 0 | 0 | 0 |
| **Gene flow from H3 into H1** | 0 | 1 | 2 | 1 | 1 | 1 | 0 | 1 | 1 | 3 |
| **Gene flow from H3 into H2** | 13 | 16 | 20 | 25 | 4 | 1 | 1 | 3 | 1 | 2 |
| **Gene flow from H4 into H1** | 0 | 0 | 0 | 0 | 1 | 0 | 0 | 0 | 0 | 0 |
| **Gene flow from H4 into H2** | 0 | 0 | 0 | 0 | 0 | 0 | 0 | 0 | 0 | 0 |
| **Gene flow between ancestor H1-H2 and H3** | 15828 | 14770 | 15226 | 14043 | 17750 | 16560 | 17029 | 15788 | 17459 | 16259 |
| **Gene flow between ancestor H1-H2 and H4** | 7 | 4 | 2 | 2 | 5 | 3 | 13 | 1 | 5 | 5 |
| **NA** | 11 | 11 | 11 | 10 | 13 | 12 | 1 | 9 | 13 | 11 |
| **None** | 5315 | 6426 | 5916 | 7126 | 3676 | 4868 | 4400 | 5644 | 3915 | 5117 |

| | | | | | | | | | | |
|---|---|---|---|---|---|---|---|---|---|---|
| **H1** | Bottlenose dolphin | Bottlenose dolphin | Pilot whale | Pilot whale | Pilot whale | Pilot whale | Pilot whale | Pilot whale | Pilot whale | Pilot whale |
| **H2** | White-sided dolphin | White-sided dolphin | White-sided dolphin | White-sided dolphin | White-sided dolphin | White-sided dolphin | Killer whale | Killer whale | Killer whale | Killer whale |
| **H3** | Beluga | Beluga | Narwhal | Narwhal | Beluga | Beluga | Narwhal | Narwhal | Beluga | Beluga |
| **H4** | Harbour | Finless | Harbour | Finless | Harbour | Finless | Harbour | Finless | Harbour | Finless |
| **Gene flow from H1 into H3** | 28 | 25 | 32 | 31 | 31 | 35 | 1 | 3 | 1 | 2 |
| **Gene flow from H1 into H4** | 0 | 0 | 0 | 0 | 0 | 0 | 1 | 0 | 0 | 0 |
| **Gene flow from H2 into H3** | 124 | 115 | 69 | 72 | 78 | 64 | 469 | 417 | 454 | 395 |
| **Gene flow from H2 into H4** | 0 | 0 | 0 | 0 | 0 | 0 | 0 | 0 | 0 | 0 |
| **Gene flow from H3 into H1** | 0 | 0 | 1 | 4 | 2 | 1 | | 1 | 2 | 1 |
| **Gene flow from H3 into H2** | 5 | 6 | 4 | 2 | 5 | 3 | 29 | 27 | 28 | 44 |
| **Gene flow from H4 into H1** | 0 | 0 | | 0 | 0 | 0 | 0 | 0 | 0 | 0 |
| **Gene flow from H4 into H2** | 0 | 0 | 0 | 0 | 0 | 0 | 0 | 0 | 0 | 0 |
| **Gene flow between ancestor H1-H2 and H3** | 16764 | 15545 | 17616 | 16460 | 16944 | 15701 | 15101 | 14030 | 14474 | 13373 |
| **Gene flow between ancestor H1-H2 and H4** | 3 | 2 | 8 | 5 | 2 | 3 | 6 | 3 | 2 | 2 |
| **NA** | 13 | 10 | 14 | 14 | 13 | 11 | 12 | 11 | 12 | 9 |
| **None** | 4607 | 5839 | 3799 | 4956 | 4467 | 5724 | 5924 | 7051 | 6570 | 7715 |

| | | | | | | | | | | |
|---|---|---|---|---|---|---|---|---|---|---|
| **H1** | Bottlenose dolphin | Bottlenose dolphin | Bottlenose dolphin | Bottlenose dolphin | Indo-Pacific Bottlenose dolphin | Indo-Pacific Bottlenose dolphin | Indo-Pacific Bottlenose dolphin | Indo-Pacific Bottlenose dolphin | Indo-Pacific Bottlenose dolphin | Indo-Pacific Bottlenose dolphin |
| **H2** | Killer whale | Killer whale | Killer whale | Killer whale | White-sided dolphin | White-sided dolphin | White-sided dolphin | White-sided dolphin | Killer whale | Killer whale |
| **H3** | Narwhal | Narwhal | Beluga | Beluga | Narwhal | Narwhal | Beluga | Beluga | Narwhal | Narwhal |
| **H4** | Harbour | Finless | Harbour | Finless | Harbour | Finless | Harbour | Finless | Harbour | Finless |
| **Gene flow from H1 into H3** | | 2 | 4 | 2 | 25 | 24 | 26 | 22 | 2 | 1 |
| **Gene flow from H1 into H4** | 0 | 0 | 0 | 0 | 0 | 0 | 0 | 0 | 0 | 0 |
| **Gene flow from H2 into H3** | 480 | 468 | 474 | 449 | 121 | 98 | 110 | 100 | 534 | 527 |
| **Gene flow from H2 into H4** | 0 | 0 | 0 | 0 | 1 | 0 | 1 | 1 | 0 | 0 |
| **Gene flow from H3 into H1** | 1 | 1 | 1 | 1 | 1 | 1 | 0 | 0 | 1 | 1 |
| **Gene flow from H3 into H2** | 29 | 28 | 27 | 48 | 5 | 3 | 6 | 6 | 26 | 32 |
| **Gene flow from H4 into H1** | 0 | 0 | 0 | 0 | 0 | 0 | 0 | 0 | 0 | 0 |
| **Gene flow from H4 into H2** | 0 | 0 | 0 | 0 | 0 | 0 | 0 | 0 | 0 | 0 |
| **Gene flow between ancestor H1-H2 and H3** | 14300 | 13209 | 13665 | 12542 | 17444 | 16309 | 16786 | 15606 | 14306 | 13255 |
| **Gene flow between ancestor H1-H2 and H4** | 5 | 3 | 1 | 1 | 5 | 6 | 2 | 2 | 6 | 3 |
| **NA** | 13 | 11 | 14 | 10 | 13 | 11 | 11 | 11 | 10 | 10 |
| **None** | 6716 | 7820 | 7359 | 8489 | 3928 | 5090 | 4602 | 5795 | 6658 | 7713 |

| | | | | | | | | | | |
|---|---|---|---|---|---|---|---|---|---|---|
| **H1** | Indo-Pacific Bottlenose dolphin | Indo-Pacific Bottlenose dolphin | Pilot whale | Pilot whale | Pilot whale | Pilot whale | Indo-Pacific Bottlenose dolphin | Indo-Pacific Bottlenose dolphin | Indo-Pacific Bottlenose dolphin | Indo-Pacific Bottlenose dolphin |
| **H2** | Killer whale | Killer whale | Indo-Pacific Bottlenose dolphin | Indo-Pacific Bottlenose dolphin | Indo-Pacific Bottlenose dolphin | Indo-Pacific Bottlenose dolphin | Bottlenose dolphin | Bottlenose dolphin | Bottlenose dolphin | Bottlenose dolphin |
| **H3** | Beluga | Beluga | Narwhal | Narwhal | Beluga | Beluga | Narwhal | Narwhal | Beluga | Beluga |
| **H4** | Harbour | Finless | Harbour | Finless | Harbour | Finless | Harbour | Finless | Harbour | Finless |
| **Gene flow from H1 into H3** | 3 | 1 | 80 | 75 | 72 | 72 | 45 | 37 | 47 | 39 |
| **Gene flow from H1 into H4** | 0 | 0 | 1 | 0 | 1 | 1 | 0 | 0 | 0 | 0 |
| **Gene flow from H2 into H3** | 575 | 475 | 32 | 33 | 36 | 29 | 30 | 40 | 38 | 37 |
| **Gene flow from H2 into H4** | 0 | 0 | 0 | 0 | 0 | 0 | 0 | 1 | 0 | 0 |
| **Gene flow from H3 into H1** | 1 | 1 | 3 | 2 | 3 | 3 | 2 | 2 | 3 | 2 |
| **Gene flow from H3 into H2** | 33 | 48 | 3 | 3 | 2 | 1 | 2 | | 5 | 2 |
| **Gene flow from H4 into H1** | 0 | 0 | 0 | 0 | 0 | 0 | 0 | 0 | 0 | 0 |
| **Gene flow from H4 into H2** | 0 | 0 | 0 | 0 | 0 | 0 | 0 | 0 | 0 | 0 |
| **Gene flow between ancestor H1-H2 and H3** | 13686 | 12641 | 17651 | 16509 | 17001 | 15778 | 17979 | 16835 | 17318 | 16117 |
| **Gene flow between ancestor H1-H2 and H4** | 1 | 2 | 7 | 3 | 1 | 2 | 8 | 3 | 4 | 3 |
| **NA** | 11 | 10 | 13 | 12 | 12 | 10 | 18 | 18 | 17 | 14 |
| **None** | 7234 | 8365 | 3753 | 4905 | 4415 | 5645 | 3459 | 4606 | 4113 | 5328 |
